# Structural assembly of the glycan-rich, chitin-reinforced adhesive of *Hydra* is coordinated by a lectin-like protein, HvAb1

**DOI:** 10.64898/2026.06.30.735459

**Authors:** Matthias Achrainer, Julia Ofer, Mario Kanetscheider, Lisa Polz, Nick Aldred, Kevin Grüner, Stefan Redl, Angelina Neumann, Anna Seybold, Bert Hobmayer, Birgit Lengerer

## Abstract

Aquatic animals deploy adhesives, in numerous essential functions, and reversibility is a key adaptation. The molecular mechanisms of reversible wet adhesion remain poorly understood. Using a model organism, the freshwater cnidarian *Hydra vulgaris,* we dissect the mechanism of molecular assembly in a secreted adhesive and uncover a glycan and protein-based architecture organized by a lectin-like protein, *Hydra vulgaris* adhesive binding protein 1 (HvAb1). We identify HvAb1 as a nonredundant organizer of the adhesive matrix, being basal-disc specific and secreted. Knockdown of HvAb1 severely impaired attachment and disrupted footprint architecture in a mosaic pattern, with only HvAb1-positive regions of the adhesive footprint retaining their normal structure. The adhesive is wheat germ agglutinin (WGA)-reactive and contains a fibrillar chitin-based sub-network, synthesized by a basal-disc-specific chitin synthase. Applying exogeneous chitinase abolished both WGA staining and *Hydra* attachment, indicating that WGA-positive components perform essential roles in adhesion. Our results therefore describe a glycan-dominated matrix, organized via a lectin-like protein (HvAb1), which is reinforced by chitin and enables reversible adhesion underwater. This establishes *Hydra* as a tractable model to better understand the principles of reversible adhesion underwater and, potentially, inform future bioinspired, sustainable adhesives.

## Introduction

Many aquatic organisms use secreted bioadhesives for locomotion, building shelters, capturing prey, and defending themselves. Although underwater adhesion has arisen multiple times in different animal lineages, and is not evolutionarily conserved *per se*, many adhesion systems appear to have co-opted the same, pre-existing physiological processes to generate adhesive secretions ^1,2^. So, despite the diversity of adhesion strategies, and adhesive components in nature, common chemical motifs often recur. This suggests the availability of a limited toolkit to taxa that may be evolving or optimising underwater adhesion^3^. For example, post-translationally modified amino acids like hydroxylated tyrosines (DOPA) and phosphorylated serines are found in the permanent adhesives of mussels, sandcastle worms and caddisfly larvae, where they enable rapid surface coupling (reviewed in ^4^). By contrast, the adhesives of temporarily adhering animals, including sea stars, sea urchins, flatworms, and limpets, are largely composed of cross-linkable extracellular matrix proteins and glycans (reviewed in ^1,5^). These convergences imply that general design rules may underpin wet adhesion across lineages. A practical route to uncovering these rules is to interrogate basal metazoans with simple body plans and powerful experimental tractability, such as *Hydra*.

*Hydra* is a small freshwater polyp belonging to the phylum Cnidaria, characterized by a simple oral-aboral body axis. It is widely recognized for its remarkable regenerative capacity and well-defined stem cell systems, which comprise two epithelial lineages (endodermal and ectodermal) and a multipotent interstitial stem cell lineage (for reviews see ^6,7^). The oral end of the animal features the head structure, including tentacles used for prey capture and a central mouth region known as the hypostome. At the aboral end lies the foot, consisting of the peduncle and basal disc, which allows the organism to attach reversibly to surfaces by secreting an adhesive substance ^8^. Ectodermal basal disc cells, which produce and store the adhesive, contain four types of *Hydra* secretory granules (HSGI-IV), with type I being the precursors for type II ^9^. Following secretion, the adhesive materials of the different granules mix and form a footprint on the substrate, characterized by a distinctive meshwork structure ^9^. HSGI and HSGII granules exhibit high nitrogen content, while HSGII-IV stain positively with Periodic Acid Schiff and lectins, indicating the presence of proteins and glycans, respectively ^8,10^. No lipids have been detected within the adhesive granules or in the secreted footprints. A lectin-binding screen has confirmed a high abundance of glycans in the adhesive and within the basal disc region, including those recognized by the lectins wheat germ agglutinin (WGA) and *Ricinus Communis* Agglutinin I (RCA) ^10^. A combination of differential RNA sequencing and *in situ* hybridization identified several candidate genes specifically expressed in the foot region, suggesting their involvement in the adhesion process ^11^. However, the molecular mechanisms and genes driving *Hydra* adhesion remain poorly understood.

In this study, we identify a glycan-binding domain-containing protein as a key functional element of *Hydra* adhesion further called HvAb1. Upon knockdown of HvAb1, animals failed to properly attach and showed a disordered adhesive structure. We demonstrate that the adhesive footprint is mainly composed of a WGA-positive material that upon loss of HvAb1 loses its characteristic structure. We demonstrate that chitin is a functional component of the adhesive meshwork and is produced by a basal disc-specific chitin synthase. Together, these findings define a process by which HvAb1 organizes a WGA-positive meshwork, including chitin, into a glycan-rich adhesive. This new understanding positions *Hydra* as a tractable model for understanding conserved principles of wet adhesion in higher taxa and may contribute to the development of biomimetic underwater glues.

## Results

### Selection of adhesion candidate proteins

To identify genes potentially involved in the adhesion of *Hydra*, we integrated data from previously published *in situ* hybridization screens, mass spectrometry-based proteomic analyses ^11^ and single-cell RNA sequencing ^12,13^. Our objective was to prioritize genes that were both components of the adhesive footprint proteome and highly expressed in basal disc cells. Using this approach, we selected fourteen proteins for further functional analyses (Fig. 1a). These include two peroxidases (Pero), two fascin-like proteins (Fas), two proteins with no predicted domains (Nd), two lectin (glycan-binding) domain proteins (Lb1 and HvAb1), four L-rhamnose lectin binding domain (Rb) proteins, one chitin-binding domain protein (Cb), and one protein containing a DOMON domain (Dom). Names reflect Interpro-predicted domain architectures, except HvAb1, which is named for its subsequently identified role as a *Hydra vulgaris* adhesive-binding protein (HvAb1). Additionally, we identified all genes expressed in the basal disc cells that would be required to produce chitin (Suppl. Fig. 1a). Single-cell RNA-Seq data revealed that one chitin synthase, (Chs) HVAEP12.G021525, was mainly expressed in the endoderm whereas the second, HVAEP2.G004025 hereafter referred to as Chs2, showed specific expression in the basal disc and, to a lesser extent, in the female germline (Suppl. Fig. 1a). In asexually reproducing polyps, Chs2 expression is restricted to the basal disc. The domain architecture of Chs2, which includes 14 transmembrane (TM) domains, an enzymatic domain and two N-terminal sterile alpha motifs (SAM) (Suppl. Fig. 1b), is characteristic of class I metazoan chitin synthases and contains all catalytic and structural motifs described within chitin synthases across species ^14^(Suppl. Fig. 2).

**Figure 1:**
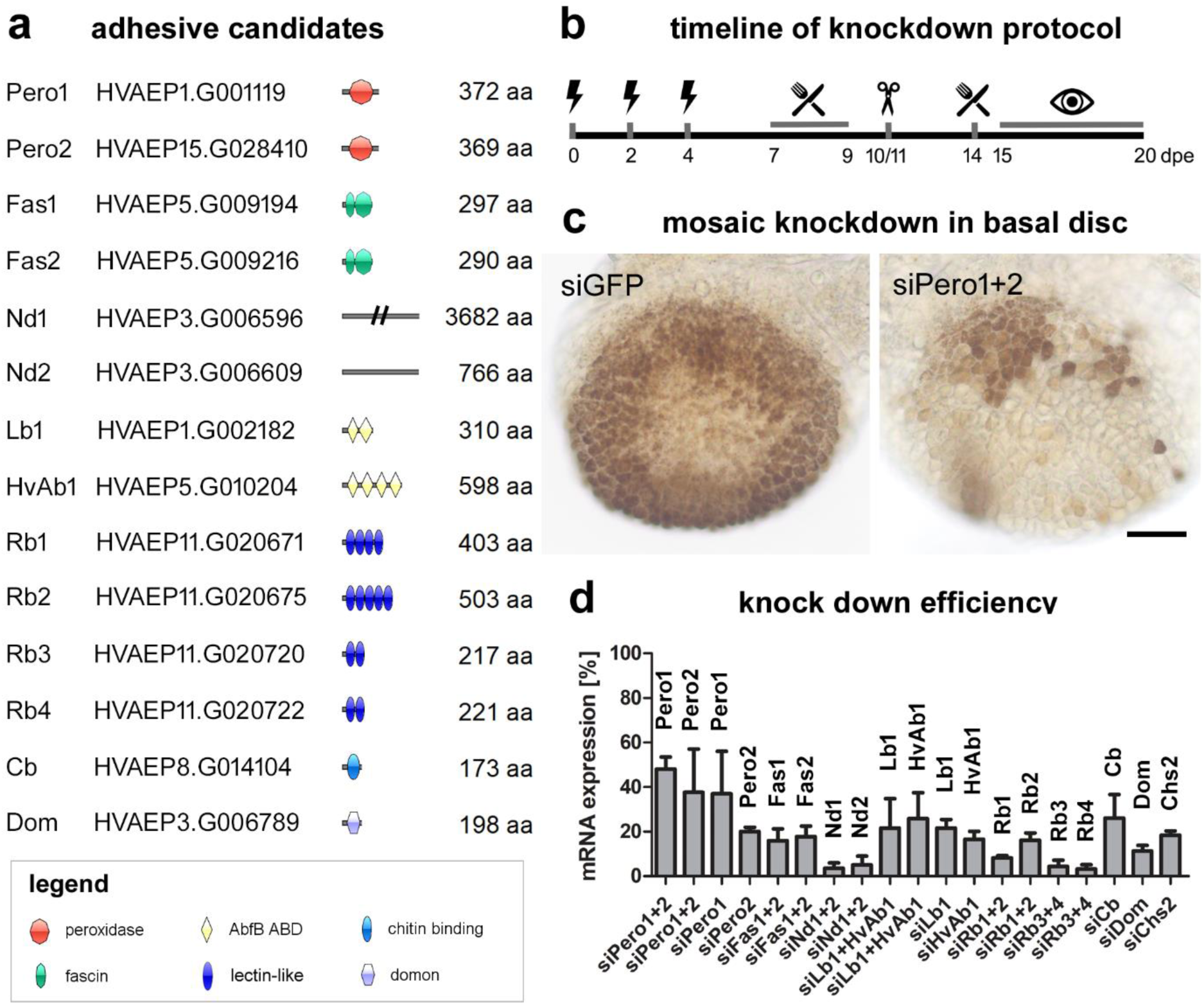
Schematic representation of siRNA-mediated knockdown targets, workflow, mosaic effect and efficiency. (a,b) Predicted functional domain organization of the secreted proteins. (c) Timeline of the siRNA knockdown protocol. (d) Mosaic knockdown pattern visualized by peroxidase (DAB) staining following control knockdown (siGFP) (n=46) and double knockdown of peroxidase genes (siPero1+2) (n=23). (e) Knockdown efficiency assessed by quantitative PCR, for double knockdowns the qPCR target is indicated on top (means of relative mRNA expression level with SD, n≥3). Scale bar in d: 100 µm.

### Knockdown of candidate genes produced a mosaic pattern in the basal disc

For functional analysis, we optimized a siRNA-mediated knockdown protocol^15^ to specifically target genes expressed in the basal disc of *Hydra*. siRNAs were delivered by electroporation three times over five days, with 24-hour recovery intervals. Each electroporation caused minor polyp shrinkage and tissue loss. After the final treatment, animals recovered for a week, then the basal disc was amputated to induce regeneration from siRNA-treated ectodermal cells. This step minimized interference from pre-existing adhesive proteins stored in secretory granules. Knockdown phenotypes were assessed 5– 9 days post-amputation (15-20 days past the first electroporation) (Fig. 1b). Histochemical staining of peroxidase activity was used to evaluate the results, with a strong signal from control animals (siGFP, Fig. 1c). The signal was markedly reduced in animals subjected to double knockdown of peroxidase genes (siPero1+2, Fig. 1c) and also revealed that the knockdown was non-uniform, leading to a mosaic signal pattern. Single knockdowns of Pero1 or Pero2 led to reduced peroxidase activity but did not abolish it (Suppl. Fig. 3), suggesting functional redundancy between the two peroxidases. Based on this observation, we hypothesized that other gene pairs with similar predicted domain motifs and structure predictions (Suppl. Fig. 4, 5), such as Fas1 and Fas2, Nd1 and Nd2, Lb1 and HvAb1, Rb1 and Rb2, Rb3 and Rb4, may also exhibit redundant functions. We performed double knockdowns targeting these candidate gene pairs to test for functional compensation. For Fas1 and Fas2, Nd1 and Nd2, Rb1 and Rb2, Rb3 and Rb4 only the double knockdown was performed using siRNAs targeting both sequences (see Suppl. Material for details). Our protocol consistently yielded high knockdown efficiency, as verified by quantitative PCR (qRT-PCR) at days 16-18 dpe (Fig. 1d). Equally high knockdown efficiencies were obtained when three different siRNAs were combined, which allowed for simultaneous knockdown of five (Lb1, Rb1, Rb2, Rb3 and Rb4) and even six genes in parallel (Fas1, Fas2, Rb1, Rb2, Rb3 and Rb4) (Suppl. Fig. 6a). Survival rates were overall around 83.4 % (SD 15.7) at 16 dpe with no significant differences between treatment groups (Suppl. Fig. 6b).

### Knockdown of HvAb1 and other adhesive genes impaired attachment abilities of *Hydra*

Specimen of *Hydra* were placed in Petri dishes containing *Hydra* medium and allowed to attach overnight. The following morning, the numbers of attached and non-attached animals were counted by two observer-blinded investigators (Fig. 2a). On average, 94.0% (SD: 4.4, n=6) of animals successfully attached in the control group and most treatment groups led to similar attachment rates. However, the knockdown of Lb1+HvAb1 showed significant reduction in the number of attached animals (35.1% SD: 7.5, n=5). Single knockdowns of Lb1 and HvAb1 revealed that this effect was caused by the knockdown of HvAb1; single knockdowns of HvAb1 led to the strongest non-adhesive phenotype with only 27.2% (SD: 12.2, n=4) of the polyps being able to attach with their basal disc. In contrast, the single knockdown of Lb1 had no effect on attachment frequency (95.8% SD: 4.6, n=4). In addition, in order to perform a quantitative analysis of attachment of polyps, we investigated removal using a calibrated water-jet system for selected treatment groups. First, the animals were placed on glass slides under steady conditions and allowed to attach. Then, remaining underwater, a water stream was applied to the animals in turn. Detachment of polyps was observed while applying increasing water flows (see Suppl. Movie 1). Wild-type and siGFP-electroporated controls showed no difference in removal, confirming the knockdown protocol itself did not affect adhesion. In contrast, siChs2, siFas1+2 and siND1+2 knockdowns resulted in moderately increased removal, while HvAb1 led to nearly complete detachment, even at the lowest wall shear stress available (Fig. 2b).

**Figure 2:**
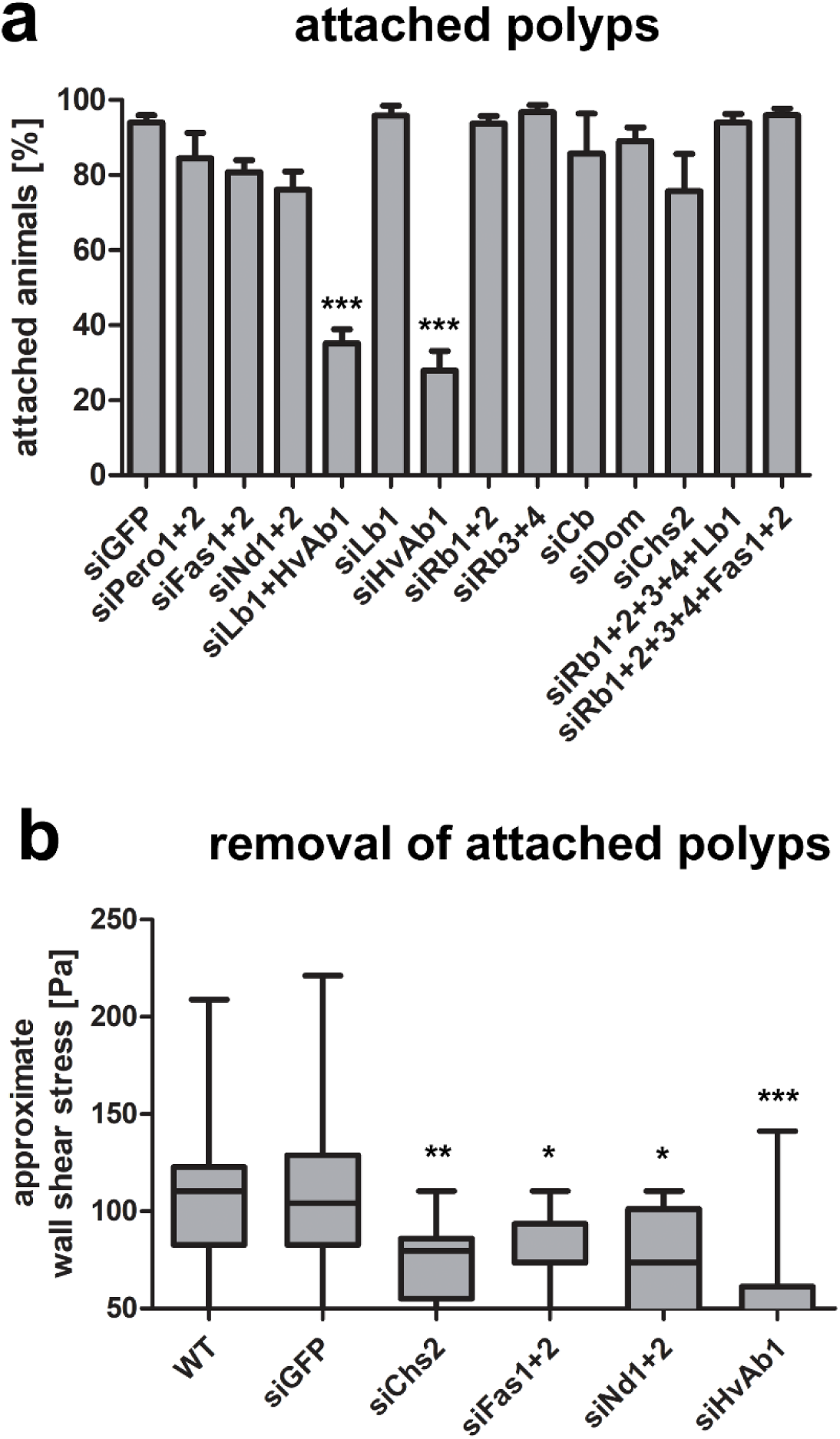
Effect of knockdown of candidate genes on attachment. (a) Number of attached animals in percentage under steady conditions. siLb1+HvAb1 and siHvAb1 are significantly lower (P<0.001) compared to siGFP (n≥3, mean with SD). (b) Attachment strength measured with water jet system, box plot indicating detached animals, line highlights median and whiskers showing min to max values. Wild-type (n=22), siGFP (n=18), siChs2 (n=19), siFas1+2 (n=16), siND1+2 (n=13), and siHvAb1 (n=22).

### Knockdown of HvAb1 disrupts the structure of the adhesive footprints

To further characterize the effects of HvAb1 knockdown on footprint formation, we generated a polyclonal antibody against an HvAb1-specific peptide. The antibody labelled only the basal disc of polyps and late-stage buds (Fig. 3a). Within basal disc cells, HvAb1 immunoreactivity did not overlap with WGA staining, indicating that the WGA-positive material and HvAb1 are not stored in the same vesicle population prior to secretion (Fig. 3b-d). In western blots of foot-enriched protein extracts, the HvAb1 antibody detected two bands at approximate 70 and 85 kDa, consistent with the predicted molecular mass of 68 kDa (Fig. 3e). We then examined the structural consequences of HvAb1 knockdown by staining polyps and their corresponding footprints with anti-HvAb1 and WGA. In control (siGFP) animals, HvAb1 localized throughout the basal disc in a pattern similar to WGA (Fig. 4a-c). In secreted footprints, HvAb1 signal was widespread and, like WGA, accumulated at the cell borders (Fig. 4d-f). Following HvAb1 knockdown, however, only a subset of basal disc cells retained detectable HvAb1 expression (Fig. 4g-i), while WGA staining remained unchanged. Animals continued to attempt attachment and deposited footprints on the substrate (Fig. 4j-l). Within these footprints, WGA labelling was widespread, however, only HvAb1-positive areas retained normal architecture with well-defined cell borders. HvAb1-negative footprints/regions lacked internal organization and discernible cell boundaries (Fig. 4j-l).

**Figure 3:**
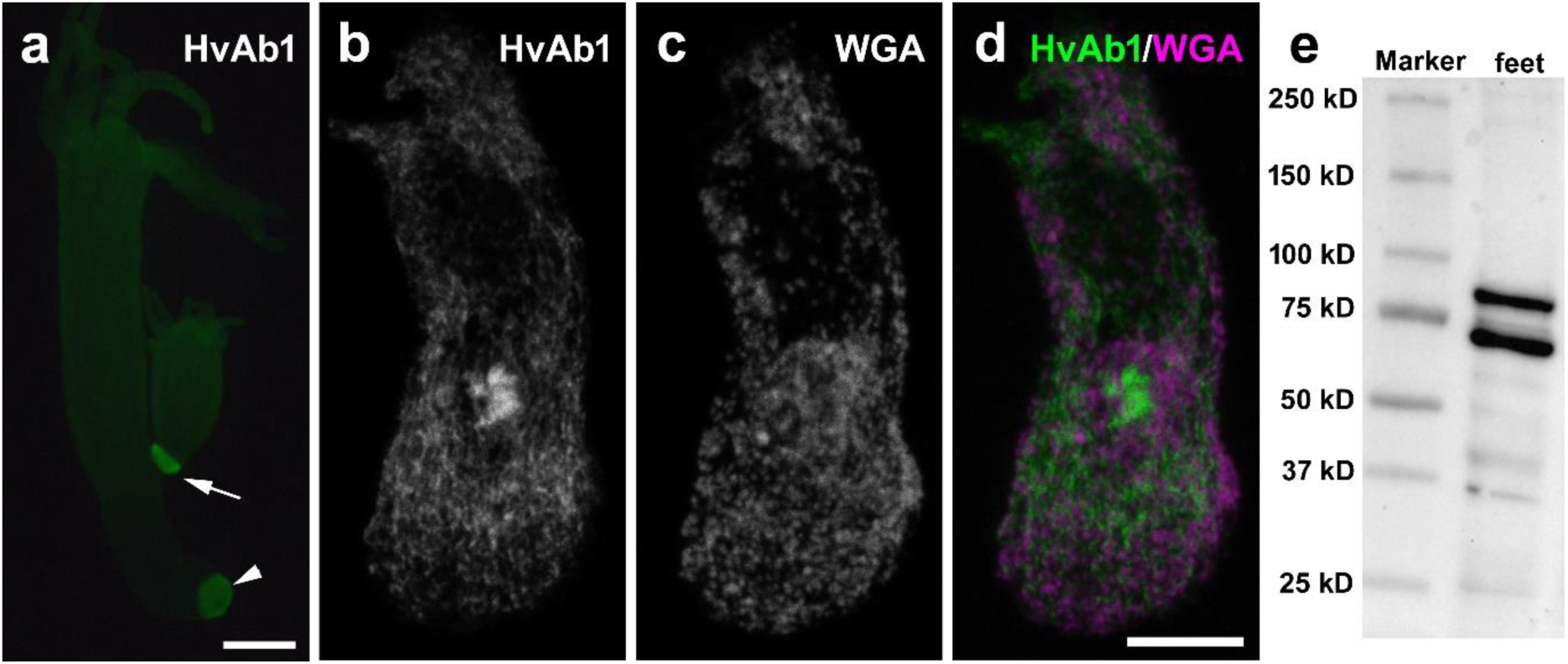
HvAb1 signal in whole mount *Hydra vulgaris* and basal disc cells and on a Western blot of basal disc enriched protein. (a) HvAb1 labeling of a polyp with a late stage bud (n=25). (b-d) Co-labelling of a basal disc cell with (b) HvAb1, (c) WGA, and (d) the merged image (n=27). (e) Western blot of basal disc enriched protein (n=6). Note the two protein bands at approximately 70 and 85 kDa. Scale bars: (a) 500 µm, (b-d) 10 µm.

**Figure 4:**
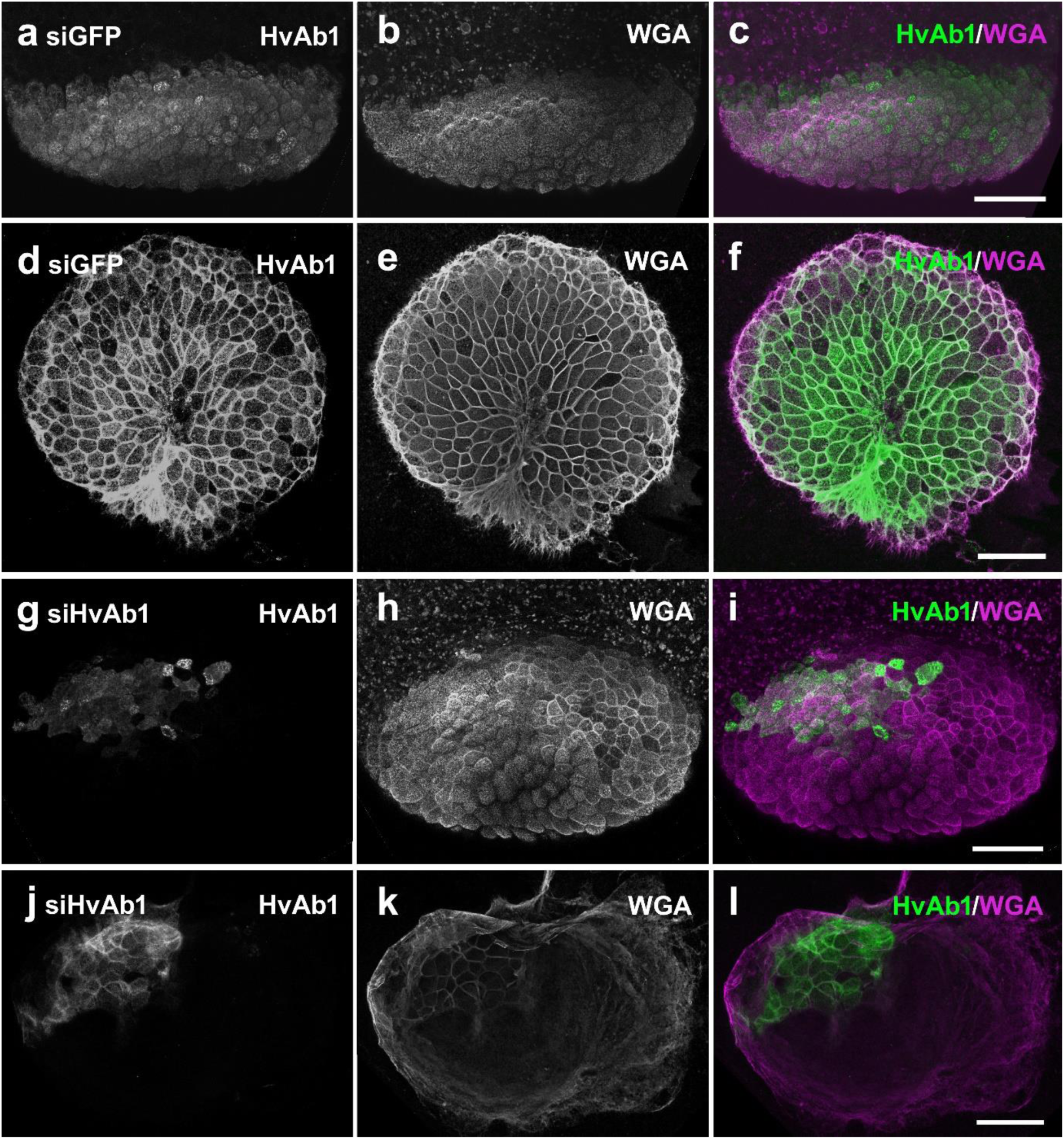
HvAb1 and WGA signals in the *Hydra vulgaris* basal disc and the corresponding footprint after siGFP (control) and siHvAb1 treatment. (a-f) siGFP treated sample: (a-c) basal disc cell co-labelled with (a) HvAb1, (b) WGA, and (c) the merged image and (d-f) the corresponding footprint co-labelled with (d) HvAb1, (e) WGA, and (f) the merged image. (g-l) siGFP treated sample: (g-i) basal disc cell co-labelled with (g) HvAb1, (h) WGA, and (i) the merged image and (j-l) the corresponding footprint co-labelled with (j) HvAb1, (k) WGA, and (l) the merged image. siGFP (n=16), siHvAb1 (n=27). Scale bars 50 µm.

### HvAb1 knockdown did not alter basal disc morphology or global gene-expression profile

Squeeze preparations of all knockdown animals showed no detectable abnormalities in basal disc morphology. Similarly, peroxidase activity, visualized using DAB staining, appeared to be normal across all siRNA treatments (Suppl. Fig. 7) except the peroxidase knockdowns (Suppl. Fig. 3). To further investigate potential ultrastructural changes, we examined siGFP (control) and siHvAb1 knockdown samples using transmission electron microscopy. No differences in the ultrastructure of basal disc cells and their granules could be observed between the samples (Suppl. Fig. 8), indicating that the cells were not visible altered by the HvAb1 knockdown. To further evaluate potential off-target effects of HvAb1 knockdown, we performed bulk RNA-seq following siRNA treatment targeting GFP (control), Lb1 (knockdown without detectable effects on adhesion), and HvAb1 (non-adhesive phenotype). As expected, both Lb1 and HvAb1 exhibited robust reductions in their mRNA levels (Suppl. Fig. 9), consistent with qRT-PCR results. Overall, only a small number of genes were differentially expressed in either knockdown relative to the control (20 in Lb1 and 17 in HvAb1; out of 11644 detected genes in both samples), indicating limited off-target effects (Suppl. Fig. 9). Analysis restricted to basal-disc specific genes revealed an upregulation of Rb4 after HvAb1 knockdown (See supplementary material for further details). Together these findings indicate that the observed non-adhesion phenotype upon HvAb1-knockdown is caused by its specific role in the adhesive formation and not due to any alterations in basal disc cell integrity.

### Chitin forms a meshwork in the *Hydra* adhesive, but makes only a minor contribution to footprint biomass

Our data suggests a role of HvAb1 as scaffolding protein organizing the structure of the adhesive, but the major components of the adhesive remained elusive. Because Chs2 knockdown reduced attachment strength and the footprints are WGA-positive, we investigated how much of the WGA signal we observed derives from chitin. We stained footprints with the chitin-specific markers CBM2A-GFP and CBP-546. Both confirmed the presence of fibrillar chitin within the *Hydra* footprint (Fig. 5a,d and Suppl. Fig. 10). Double staining of CBM2A-GFP and the lectin WGA showed that the WGA signal spans the entire footprint, whereas chitin is confined to fibrillar structures. This indicated abundant WGA-reactive components that are not chitin and which are currently undefined (Fig. 5a-f). In Chs2 knockdowns, CBM2A-GFP staining was strongly reduced and showed a mosaic pattern, confirming impaired chitin synthesis (Fig. 5g). Yet, the overall footprint architecture and strong WGA labelling persisted (Fig. 5h-i). Chs2 knockdown did not alter WGA or HvAb1 patterns, indicating that chitin is not required for adhesive organisation (Suppl. Fig. 11). By contrast, HvAb1 knockdowns disrupted both the WGA-positive material and chitin (Suppl. Fig. 12). All other knockdowns showed normal adhesive footprint structure (Suppl. Fig. 13-15). Like the HvAb1 knockdown, Chs2 knockdown had no effect on the basal disc and granule ultrastructure (Suppl. Fig. 16).

**Figure 5:**
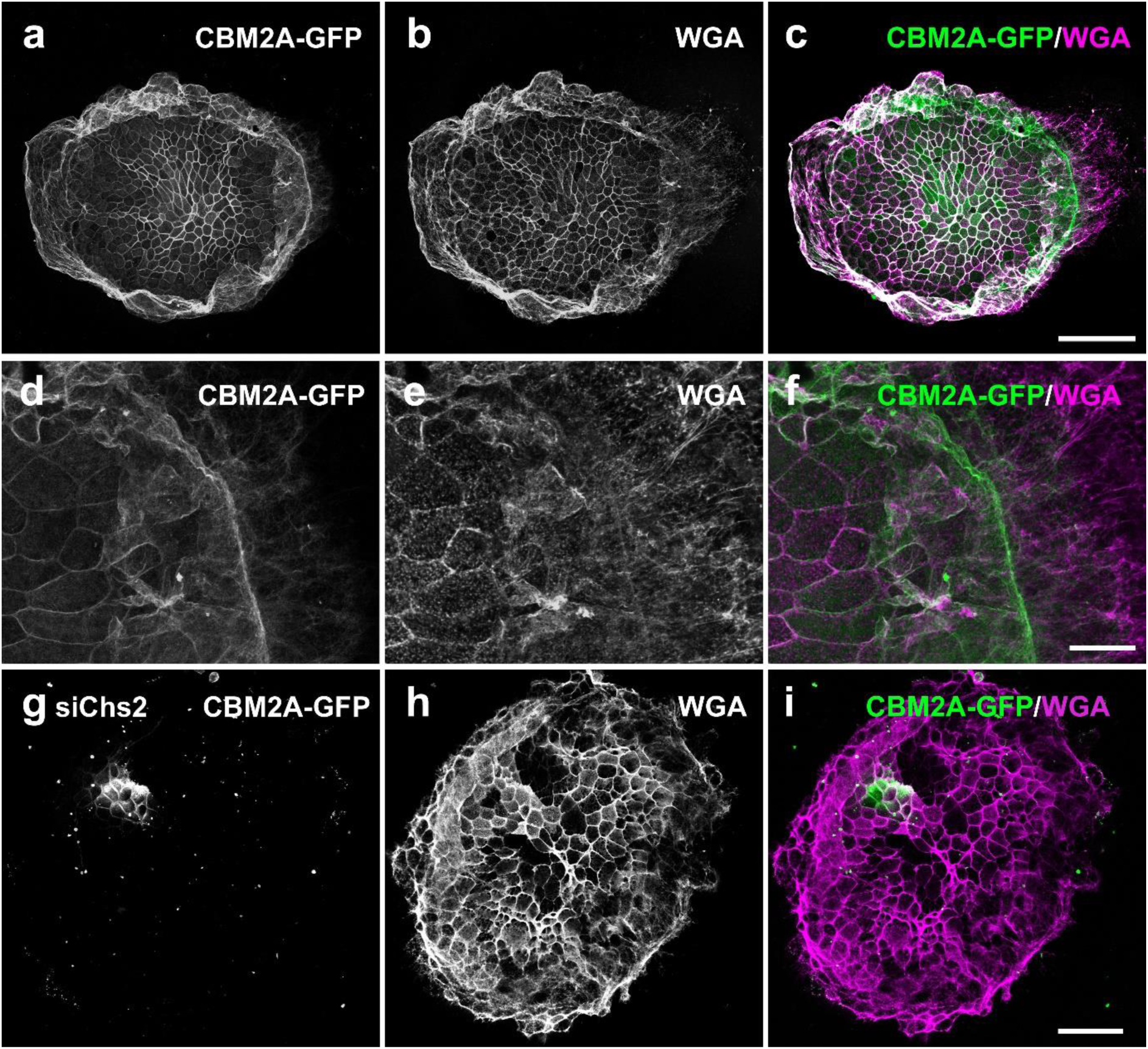
Presence and functional role of chitin in the adhesive footprint of *Hydra*. (a-f) Double labeling of chitin using CBM2A-GFP and WGA in *Hydra* adhesive footprints (n=40). Note the grainy appearance of WGA-positive but CBM2A-GFP–negative regions, suggesting the presence of non-chitinous glycoconjugates. (e-g) chitin (CBM2A-GFP) and WGA signal in a footprint secreted from a siChs2 knockdown animal (n=16). Scale bars: (c,i) 50 µm, (f) 20 µm.

### Chitinase inhibits attachment in a dose-dependent manner and degrades the adhesive meshwork

To test the functional contribution of WGA-positive components to adhesion, we incubated live *Hydra* polyps with graded concentrations of chitinase. Notably, the used chitinase also acts as N-acetyl-glucosaminidase, degrading GlcNAc disaccharides. Chitinase caused a dose-dependent reduction in attachment (Fig. 6a, Suppl. Fig. 17a-f). To evaluate structural effects on the adhesive, footprints were treated with cellulase, active chitinase, or heat-inactivated chitinase and then stained with WGA. Neither cellulase nor heat-inactivated chitinase affected the footprint architecture or WGA signal (Fig. 6b). In contrast, active chitinase treatment abolished staining with WGA (Fig. 6c), indicating enzymatic degradation of the adhesive. Consistent with these findings scanning electron microscopy of fixed polyps showed that cellulase-treated basal discs retained the characteristic fibrous meshwork of the adhesive layer (Suppl. Fig. 17g), whereas this structure was lost after chitinase treatment, confirming degradation of the adhesive matrix (Suppl. Fig. 17h). In summary, these findings indicated that GlcNAc components in the adhesive are required for adhesive structure and attachment.

**Fig. 6:**
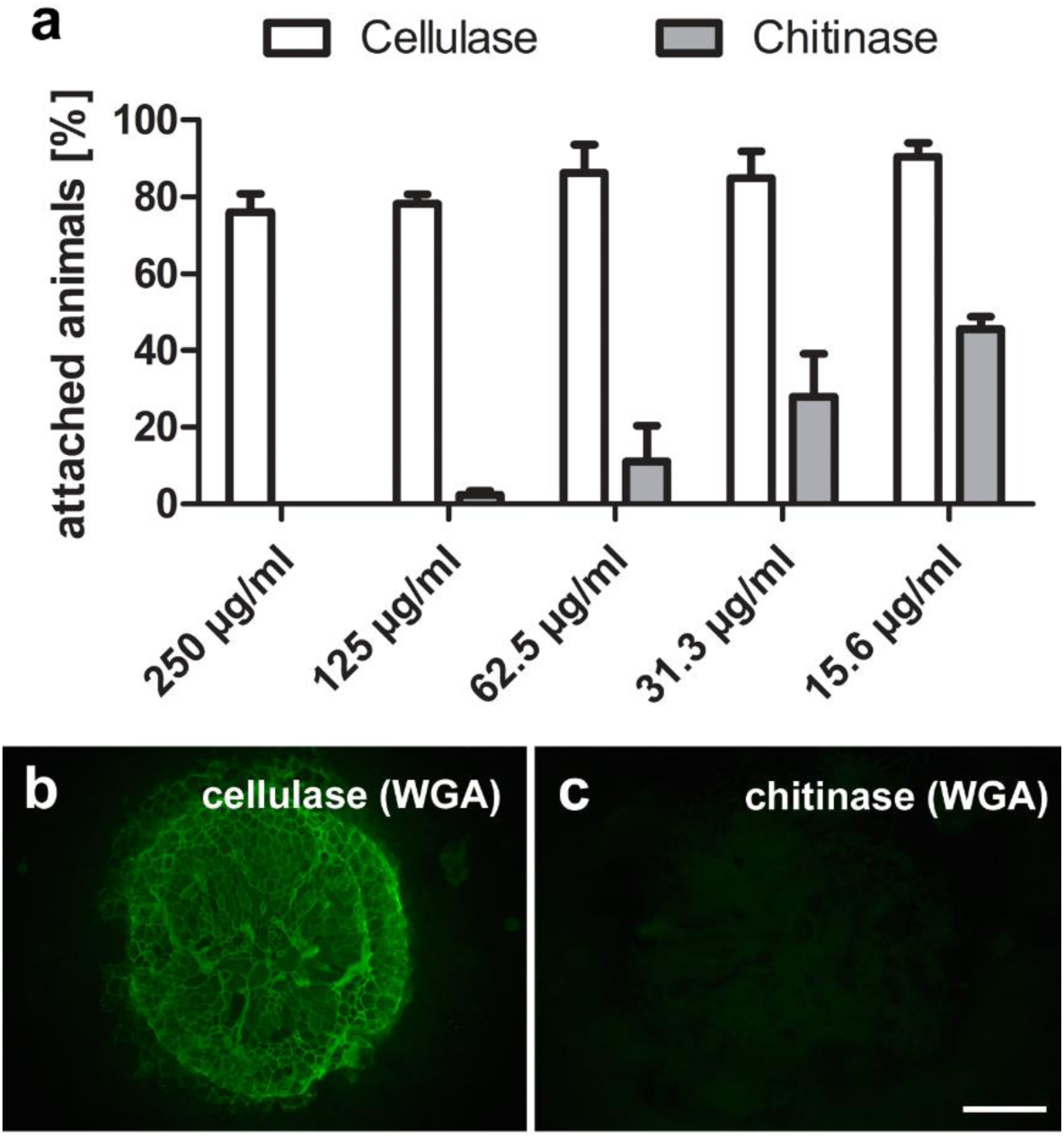
Chitinase treatment impairs attachment of living *Hydra vulgaris* and diminishes secreted footprints. (a) Quantification of attached animals following incubation with decreasing concentrations of cellulase and chitinase, error bars represent standard deviation (n=3, with 15 individuals each, P<0.001). (b,c) WGA staining of footprints after enzymatic treatment with (c) cellulase (n=35) or (c) chitinase (n=35). Scale bars 100 µm.

## Discussion

### HvAb1 is a nonredundant organizer of a glycan-based adhesive in *Hydra*

From a shortlist of 14 candidates, HvAb1 emerged as a key determinant of attachment. HvAb1 is a secreted glycan-binding protein that is restricted to basal disc cells and is deposited into footprints, consistent with a direct role in adhesion. Within cells, HvAb1 and WGA-reactive cargo do not colocalize, suggesting separate packaging and coordinated extracellular assembly. In secreted footprints, both HvAb1 and WGA signals are widespread and enriched at apparent cell borders.

Functionally, HvAb1 knockdowns produced a clear weakening effect on attachment. Animals attempted to attach but failed to remain adhered. Importantly, footprints were still deposited, but regions lacking HvAb1 were disorganized and lacked the characteristic internal architecture. HvAb1-positive patches retained normal structure. This mosaic phenotype provided an internal control that localized HvAb1’s action in the matrix rather than having a role upstream (e.g. changes in cell identity, secretory capacity, or ultrastructure, all of which appeared normal). Knockdown of other glycan-binding proteins, including combined knockdowns of up to six, did not abolish attachment or alter footprint morphology. However, simultaneous knockdown of Fas1 and Fas2 caused a modest reduction in attachment strength, suggesting that other glycan-binding proteins also contribute to internal stabilization of the adhesive. Only HvAb1, however, serves a nonredundant role. Bulk RNA-seq confirmed HvAb1 silencing with minimal off-target transcriptional changes, further supporting a primary material-level role. Modest upregulation of Rb4 suggest a compensatory response that was insufficient to restore function, further highlighting the essential role of HvAb1.

### Functional roles of the WGA-positive matrix in adhesion

Exogenous chitinase produced a dose-dependent loss of attachment and strongly reduced adhesive material on both basal discs and substrates. Chitinase abolished WGA staining, consistent with the enzymatic disruption of GlcNAc-containing components beyond chitin. The severity of the chitinase phenotype, together with the broad distribution of WGA reactivity across footprints, highlights the WGA-positive matrix as essential for adhesion. RNAi-mediated knockdown and enzymatic perturbation dissociated the roles of the WGA-reactive footprint material and chitin. Chs2 knockdown yielded a moderate reduction in attachment strength while preserving overall footprint architecture, indicating that the WGA-positive network can form independently of chitin. Chitin, while forming a fibrillar sub-network that enhances cohesion, is therefore supportive rather than required for the establishment of footprint architecture.

### A working model for the *Hydra* adhesive matrix

Our data support a model in which the *Hydra* basal disc secretes a glycan-rich adhesive dominated by WGA-reactive materials that are structurally organized by HvAb1 and reinforced by chitin fibrils (Fig. 7):

- Packaging and secretion: HvAb1 and WGA-positive cargoes are stored in distinct granule populations and co-deposited onto the substrate. Upon secretion, WGA-reactive material spreads broadly and defines the footprint. HvAb1 also distributes widely, consistent with a role in matrix assembly.
- Matrix assembly and architecture: HvAb1 crosslinks or templates interactions among WGA-reactive glycoconjugates and potentially links the adhesive to the cell surface of the animals, establishing adhesive architecture and the characteristic cell-border pattern in the footprint. Loss of HvAb1 reduces adhesive organization, yielding structurally deficient footprints and failure to sustain attachment. Nevertheless, footprints still form and adsorb to the substrate in HvAb1 knockdown animals, indicating that HvAb1 is dispensable for interfacial binding with a substrate. HvAb1 is predicted to encode a protein with four glycan-binding sites. We propose that these sites potentially crosslink WGA-positive components, as well as, chitin fibres to generate a functional adhesive.
- Mechanical reinforcement by chitin: Two independent probes revealed a fibrillar chitin sub-network embedded within the footprint, while most of the footprint remained non-chitinous but WGA-reactive. Knockdown of the basal-disc chitin synthase Chs2 diminished chitin and reduced attachment strength without abolishing overall footprint architecture or WGA signal.

**Fig. 7:**
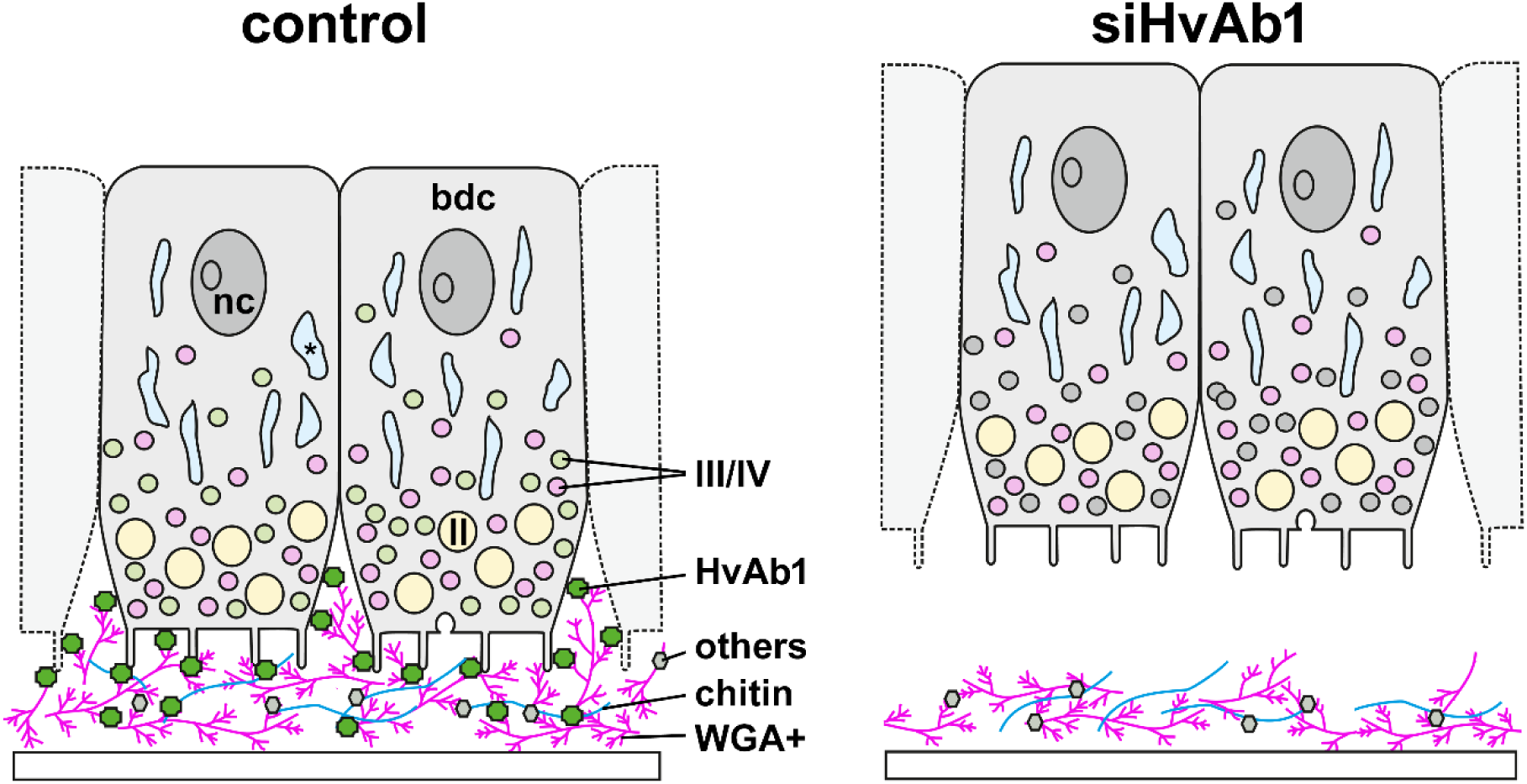
Schematic representation of basal disc cells and the secreted adhesive in control and siHvAb1 knockdown animals. Under normal conditions the basal disc cells stay attached to the substrate (white bar at the bottom). The WGA-positive material adsorbs to the substrate and forms the bulk of the adhesive material (drawn in magenta) and is modulated and attached to the basal disc cells by HvAb1 (drawn in green). Adhesive material accumulates at the basal disc cell borders. Chitinous fibres (drawn in blue) and other secreted proteins intermingle and increase attachment strength. In the absence of HvAb1, the adhesive adsorbs to the substrate, but the cells fail to stay attached and no cell borders can be distinguished in the secreted adhesive footprint. Abbreviations: **II** adhesive granules type II, **III/IV** adhesive granules type III/IV, **bdc** basal disc cell, **nc** nucleus, asterisk highlights water vacuoles.

### Context within other aquatic adhesives

*Hydra* is a freshwater polyp within Cnidaria, a lineage that evolved underwater adhesion early in metazoan history ^16^. Despite the extensive use of cnidarians as model organisms and increasingly rich experimental toolkit, relatively few studies have addressed their adhesive systems. Work to date has focused mainly on *Hydra vulgaris* ^9,11^ and the marine anemone *Exaiptasia pallida* ^16,17^. Although both are solitary polyps, their basal discs differ markedly in ultrastructure ^9,16^. *E. pallida* secretions are WGA-positive but chitin-negative, indicating a glycan-rich adhesive with distinct composition. Several proteins upregulated in the basal disc appear conserved between *Hydra* and *E. pallida*, including Rb3, suggesting shared modules. However, overall organization differs between the organisms, and functional validation has not been performed in *E. pallida*.

Aquatic adhesives are typically composite secretions of proteins and glycans, and *Hydra* appears to be no exception. Because proteins are more accessible analytically, they have been studied more thoroughly (reviewed in ^18–20^). In marine, temporarily adhering animals, adhesive secretions are dominated by large, multimodular, fibre-forming proteins ^17,21–27^. Although not conserved across species, these proteins share key features, including biased amino acid composition, extensive repeats and diverse conserved functional domains like van Willebrand factor domains, EGF-like domains and glycan-binding (lectin-like) domains ^1^. Some of these marine proteins also retain function under freshwater conditions ^23,28^. In *Hydra*, comparably large, fibril-forming proteins have not been detected in the footprint proteome ^29^. Instead, the adhesive features multiple shorter proteins with glycan-binding domains and is rich in broadly WGA-reactive glycoconjugates including a chitin sub-network. Many marine adhesives likewise contain abundant WGA-positive components ^16,30–32^, yet the specific roles of glycans remain less explored than those of proteins. In mussels, proteins with EGF-like domains bind N-acetylglucosamine and thereby contribute to robust underwater adhesion ^33^. The authors argue that domains annotated as EGF-like in adhesive proteins might be frequently glycan-binding in function, and such modules are widespread across diverse adhesive repertoires ^17,21–27^. Similar protein-glycan interactions are often implicated in aquatic adhesives, but remain incompletely characterized across taxa. *Hydra* offers a complementary freshwater model with *in vivo* tractability.

### The role of chitin within the adhesive

In cnidarians, chitin has been recognized as a key component of the rigid exoskeletons and holdfasts of benthic Medusozoa ^34^. Previous studies have also identified chitin synthases in cnidarians that lack exoskeletons ^14,35^ and localized chitin to the *Hydra* foot ^35^. But the functional role of chitin in *Hydra* was previously unknown. We show that chitin forms flexible fibres in the secreted adhesive, enhancing cohesive strength, and is synthesized by a basal-disc-specific chitin synthase. To our knowledge, chitin as a component of an aquatic adhesive has only been described previously in the hardening cement of barnacles ^32^. Its role there is perhaps not surprising given the importance of chitin in the formation of arthropod cuticle, however *Hydra* is a soft-bodied organism, requiring an adhesive that remains flexible and supports voluntarily detachment – a key functional distinction from permanently adhering organisms like barnacles.

### Outlook and future perspectives

Our data underscore the importance of WGA-positive components in the adhesive, although their molecular identities remain unresolved. A subset of these WGA-positive signals corresponds to chitinous fibres, which enhance attachment strength. Chitin and chitosan are sustainable interfacial materials: biodegradable, biocompatible, essentially non-toxic and endowed with attractive physicochemical properties. Yet their use in tissue-specific or cell-adhesion applications remains limited. Here, we identify HvAb1, a *Hydra* lectin-like protein that operates in fully aquatic conditions and potentially binds both N-acetylglucosamine-containing glycans and chitin fibres. If its binding specificity can be experimentally confirmed, HvAb1 could inspire a molecular coupling agent that organizes glycan-rich matrices under wet conditions and offer design principles for next-generation, reversible and sustainable adhesive biomaterials.

## Materials and Methods

### *Hydra* culture

All experiments were carried out with individuals of *Hydra vulgaris* strain AEP, which were bred and kept in mass cultures at the Institute of Zoology, University of Innsbruck. *Hydra* cultures were kept in growth chambers at 18 °C in *Hydra* culture medium and fed five times per week with *Artemia nauplii.* Before any experiment, the animals were starved for 24 h.

### siRNA design and selection

The siRNA sequence targeting GFP, used as a control, was obtained from ^36^. Candidate-specific siRNAs were designed using the online tools siDirect v2.1 ^37^ and RNAxs ^38^ (sequences listed in Suppl. Table 1). Off-target effects were minimized by performing BLAST searches against the Hydra AEP genome using the *Hydra* AEP Genome Browser hosted by the National Human Genome Research Institute. siRNA candidates exhibiting fewer than four mismatches with any off-target sequences were excluded from further analysis. Final siRNA sequences used in the study are provided in Suppl. table 1. siRNAs were ordered from Microsynth with an rUrU 5’ and 3’ overhang.

### siRNA mediated knockdown procedure and phenotype characterization

For each treatment, 20 bud-free *Hydra* polyps were selected and transferred to a 4 mm electroporation cuvette. Polyps were washed with RNase-free water. Prior to the first electroporation, five washing steps were performed; only two washing steps were conducted before the second and third electroporation. After washing, RNase-free water was replaced with 200 µl of 4 µM siRNA solution. For single knockdowns one siRNA was used at an end concentration of 4 µM. For double and triple knockdowns, the concentration of siRNAs was lowered to a concentration of 2 µM (for details on the siRNAs and concentrations used in the various knockdown combinations see Suppl. Material). Polyps were electroporated with a square pulse of 250 V and 25 ms in a Gene Pulser X cell^TM^ (Biorad). Immediately after electroporation precooled recovery medium ^15^ was added and the polyps were gently transferred to a Petri dish prefilled with cooled recovery medium. After 24 hours the polyps were transferred from recovery medium to *Hydra* medium. Electroporation was performed on 0, 2, and 4 days past electroporation (dpe). Animals were fed four times (7, 8, 9 and 14 dpe). As siRNA knockdown was less efficient in the basal disc, the feet of the polyps were amputated at 10 or 11 dpe. The knockdown phenotypes were investigated after complete basal disc regeneration from 15 to 20 dpe. Number of overall and attached animals were counted in Petri dish filled with *Hydra* medium by two observer-blinded investigators.

### Evaluation of knock down efficiency

Knock down efficiency was evaluated with qRT-PCR. Primers were designed with primer3 with standard settings and are listed in (Suppl. Table 2) First, the RNA of the cohort of knockdown animals was isolated using the Monarch® Total Miniprep Kit (New England BioLabs) according to the manufacturers protocol. Then, the RNA concentrations were measured with a nanodrop. For all samples of one knockdown experiment, either 700 ng or 1 µg of template RNA was used for the reverse transcription to cDNA with the SuperScriptTM IV First-Strand Synthesis System (Themo Fisher Scientific) or the LunaScript® RT SuperMix (New England BioLabs). For the qRT-PCRs the Luna® Universal qRT-PCR Master Mix (New England BioLabs) was used according to the manufacturers protocol. The qRT-PCR was performed with the QuantStudioTM 3 System. The data was analysed focussing on the CT-values (number of cycles needed to detect the gene) ^39^. In the first step, the CT-values of the genes of interest were normalised against the expressions of two housekeeping genes (EF 1α an RP 15). After this, the relative expression patterns of the genes were calculated with N=2^(-ΔΔCT). For each treatment, at least three biological replicates were made. For statistical analysis, data was evaluated in Prism through one-way ANOVA and Bonferroni’s multiple comparisons post-test.

### Staining of peroxidase activity in the basal disc

Polyps were relaxed in 2 % urethane in *Hydra* medium for 4 min and fixed for 1 h in 4 % PFA in PBS at RT. After washing in PBS-T, staining was performed using 1 ml DAB substrate solution and 1 drop of Chromogenic substrate (Sigma Aldrich). Staining was performed under visual observation and stopped when controls were stained dark brown. To stop the reaction, polyps were washed three times in PBS-T and mounted in Mowiol (Roche).

### Removal under hydrodynamic shear

To analyse the attachment strength of *Hydra* polyps under standardised conditions, water jet measurements were performed at the University of Essex (UK). These experiments used a purpose-built and calibrated, robotic water-jetting system (Suppl. Fig. 18). The equipment was designed by Dr. Nick Aldred and built by Advanced Analysis & Integration Ltd., Manchester (UK). Unlike similar instruments presented previously ^40,41^, this system allowed for the removal of adhered organisms using a jet of water while the animals remain totally immersed. It comprises an acrylic tank with a horizontal slide-holding rack, above which a gantry system raster-scans a vertical nozzle assembly over the slides, with organisms attached, at a defined stand-off distance. The slide holder can accommodate up to 12 slides simultaneously. Water is recirculated through the tank and nozzle using a diaphragm pump, and the scan pattern is loaded into the on-board computer using a graphical interface on a Wi-Fi enabled tablet. Impact pressure is controlled using a mechanical purge valve and pump-pressure gauge. Pump pressure on the gauge corresponds to impact pressure according to the calibration (Suppl. Fig. 19). For the experiments described here, we used a stand-off distance of 10mm and a 1.5mm nozzle diameter.

Hydra polyps were allowed to attach to glass slides for 15 minutes while in the slide holder, immersed in the water tank. Polyps were removed from the edges of the slides prior to experiments and the system was programmed to run with the water jet impacting the slide edges, around 1 cm away from attached polyps. In this way, polyps were never directly hit by the jet from above and, instead, experienced shear forces from water spreading away from the impact point, parallel to the surface (Suppl. Fig. 20).

Because the polyps were within the so-called impingement region radiating away from the point of impact, the flow they experienced at the ‘viscous similar’ point was expected to be laminar and the wall shear stress could be approximated using the equation 40:

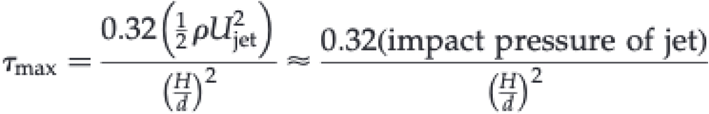

Where impact pressure is measured in Pa, *H* is the nozzle stand-off distance and *d* is the nozzle diameter. It should be noted that this method provides approximate values for wall shear stress and not precise values. Raw data for impact pressure are provided in supplemental data. However estimated wall shear stress is used here for its intuitive meaning; for context, the lowest reported wall shear stress of 49 Pa, estimated at a gauge pressure of 0.2 Bar, would be roughly equivalent to the shear stress on the hull of a ship travelling at around 10 knots ^40^. Experiments began at a minimum impact pressure of 49 Pa and repeated through increasing pressure cycles until all polyps were removed (see supplemental movie), with number remaining recorded after each pressure increase and an average removal pressure for each treatment estimated at the end.

### Antibody and WGA staining of Hydra polyps, basal disc cells, and footprints

The HvAb1-specific peptide KNNVNPDDASMDETC was selected for polyclonal antibody production in rabbits. The polyclonal antibodies were isolated from the crude serum by affinity purification using the synthetic peptide (Genscript, USA). For basal disc cell staining the basal disc were macerated as described in ^10^ and afterwards the cells were stained on glass slides as described below. For antibody staining after siRNA treatment, the polyps were individualized at day 15 dpe and kept in uncoated ibidi 8-well slides (Fisher Scientific) overnight. The footprints and individual polyps were simultaneously labelled within the ibidi wells, allowing to assign the footprints to the individual animals. The animals were relaxed in 2% Urethan in *Hydra* medium for approximately 4 min and afterwards fixed for 4 hours in Ladovsky fixative (ethanol:formamid:acetic acid:distilled water— 50:10:4:40) at RT. After several washing sets in in Tris buffered saline (25 mmol l-1 Tris, 125 mmol l-1 NaCl, pH 8.) containing 0.05% (v/v) Triton (TBS-T) the samples were blocked in TBS-T containing 3% (w/v) bovine serum albumin (BSA-T) for at least 1 h at RT. The anti-HvAb1 antibody was diluted 1:500 in BSA-T containing 5 µg/ml biotinylated WGA and added to the samples overnight at 4°C. After several washing steps in TBS-T a 1:700 diluted goat-anti-rabbit-Alexa488 antibody (Abcam) and 1:500 diluted Streptavidin-Texas Red (Vector) in BSA-T were applied for 1 hour at RT. After several washing steps in TBS-T, the individual polyps were transferred onto a glass slide and both the polyps and footprints were mounted in Vectashield® (Vector). All samples were analysed with a Leica DM5000 microscope and a Leica Sp5 confocal microscope.

### HvAb1 Western blot of basal disc enriched protein

150 Hydra feet were dissected and homogenized in 800 µl lysis buffer with protease inhibitors (Halt Protease Inhibitor Cocktail, Fisher Scientific) by pipetting. The homogenised tissue was transferred to a new precooled 1.5 microcentrifuge tube and centrifuged at maximum speed for 10 min at 4°C. Supernatant was transferred to a new tube, with the pellet being discarded. Protein concentration was quantified using the Pierce 660 nm Protein Assay (Thermo Scientific) and BSA standards (Thermo Scientific) and measured on a Nanodrop2000c. Proteins were diluted in 4XLaemmli buffer (Bio-Rad) and 5 µl were loaded onto a Criterion TGX stain-free gel (Bio-Rad). Precision Plus Protein™ All Blue Standard (Bio-Rad) was used as a marker. The gel was run with a Bio-Rad PowerPac using the Tris-glycine program 110 V for 60 min. Semi-dry blotting was performed as with a Bio-rad Trans-Blot Turbo system to a PVDF membrane with 25V and 1A for 30 min.

Afterwards the membrane was washed in 1xTBS-T and blocked with 5% milk powder in TBST overnight at 4°C. Then the membrane was incubated in 1:5000 diluted anti-HvAb1 antibody in 5% milk powder in TBST for 1 hour at RT, followed by several washing steps in 1xTBS-T. Then the membrane was incubated in 1:10000 diluted secondary antibody (mouse-anti-rabbit-HRP, Abcam) for one hour at RT., followed by several washing steps in 1xTBST. For detection, the ECL Select^TM^ Western Blotting Detection Reagent (GE Healthcare) was used. After incubation for five minutes the membrane was exposed in a ChemiDoc XRS (Bio-Rad) for image capture.

### Chitin and WGA staining of *Hydra* footprints

Footprints were collected by placing polyps on microscope glass slides, submerged in *Hydra* medium, or in 8-well ibidi slides. After at least one hour up to overnight, the animals were gently detached using a glass pipet and the slides fixed in 4 % paraformaldehyde (PFA) in phosphate buffered saline (PBS) for 30 min at room temperature (RT). Samples were washed several times in Tris-buffered saline (TBS, pH 8.0) supplemented with 5 mM CaCl2 and 0.1% Triton X (TBS-T). Unspecific background staining was blocked by pre-incubation in TBS-T containing 3% (w/v) bovine serum albumin (BSA) for 1 h at RT. Chitin-binding modules were diluted in 3 % BSA in TBS-T (CBM2A-GFP: 1:25 diluted; CBD-546 1:20 diluted) and applied overnight at 4 °C. After washing with TBS-T, biotinylated WGA (Vector) 10 µg/mL diluted in 3 % BSA in TBS-T was added for 1 h at RT. After washing with TBS-T, samples were incubated in 1:500 diluted Texasred-conjugated-streptavidin (Vector) in 3 % BSA in TBS-T for 1 h at RT. After washing with TBS-T, the samples were mounted in Vectashield. For WGA single staining, WGA conjugated with fluorescein (Vector) was used at a concentration of 10 µg/mL diluted in 3 % BSA in TBS-T for 1 h at RT. The samples were analyzed with a Leica DM5000 microscope or with a Leica SP5 II confocal scanning microscope.

### Chitinase and cellulase treatment

For chitinase and cellulase treatment of living animals, chitinase and cellulase were diluted in *Hydra medium* and the animals submerged in the different concentrations for 24 hours, then the attached animals were counted by at least two investigators (observer-blinded). For chitinase and cellulase treatment of footprints, footprints were collected on clean glass slides. After fixation in 4 % PFA in PBS and several washes in TBS-T. Chitinase (from *Streptomyces griseus*, Sigma Aldrich) or Cellulase (from *Aspergillus niger,* Sigma Aldrich) 0.5 mg/mL in TBS pH 5.5 was applied overnight at 37 °C. After several washing steps with TBS-T, footprint staining was performed with WGA conjugated with fluorescein (Vector) 10 µg/mL.

## Data accessibility

Sequencing data have been deposited in NCBI Gene Expression Omnibus with the accession: GSE338599.

## Authors’ contributions

Conceptualization, B.L.; Methodology, M.A., J.O., M.K., K.G., N.A., S.R., A.N., A.S. and B.L.; Formal Analysis, M.A, J.O., M.K., N.A., B.H. and B.L.; Writing – Original Draft Preparation, M.A. and B.L.; Writing – Review & Editing, M.A., N.A, B.H. and B.L.; Visualization, M.A. and B.L.; Supervision, B.L.; Funding Acquisition, B.L.

## Declaration of Competing Interest

The authors declare that they have no competing financial interests or personal relationships that could have influenced the work reported in this paper.

## Funding

B.L. is funded by an ESPRIT grant of the Austrian Science Fund (FWF): [ESP 15]. BL, and N.A. were members of the COST Action “European Network of Bioadhesion Expertise” (CA15216). N.A. is grateful for support by the Office of Naval Research (N00014-23-1-2177-P00001 & N00014-25-1-2529) and ONR Global, who funded the development of the waterjet apparatus (HR0011-23-2-0008).

## Acknowledgements

The authors thank Natalie Kolb for excellent *Hydra* culture maintenance and Bianca Horrer and Marian Hirschfeld for technical assistance.

## Notes

### Competing Interest Statement

The authors have declared no competing interest.

### Summary of Updates

Corrected a typographical error in the abstract and added the sequencing data accession number.

