## Supplementary material for "Structural assembly of the glycan-rich, chitin-reinforced adhesive of *Hydra* is coordinated by a lectin-like protein, HvAb1"

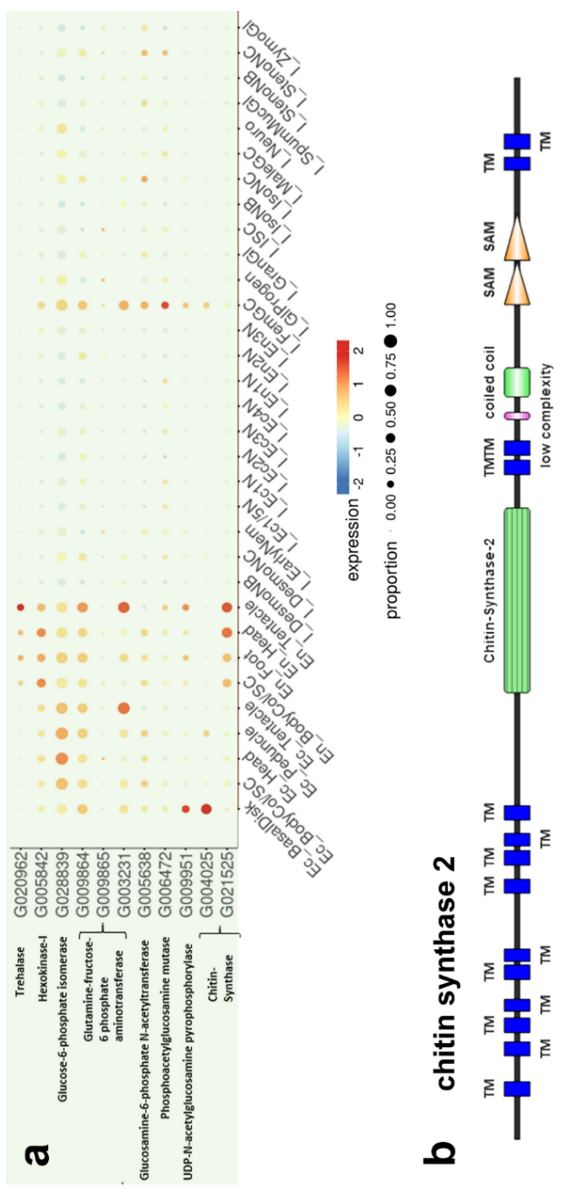


**Suppl. Fig. 1: Expression profile of the genes required for chitin synthesis and protein domain structure of the basal disc expressed chitin synthase 2 (Chs2).** (a) Expression profiles of genes involved in the chitin synthesis according to the single cell browser of the National Human Genome Research Institute. Size of the dots represent the percentage of cells expressing the gene, while the colour reflects the average expression level of the gene in each cell. (b) Predicted functional domain organization of Chs2.

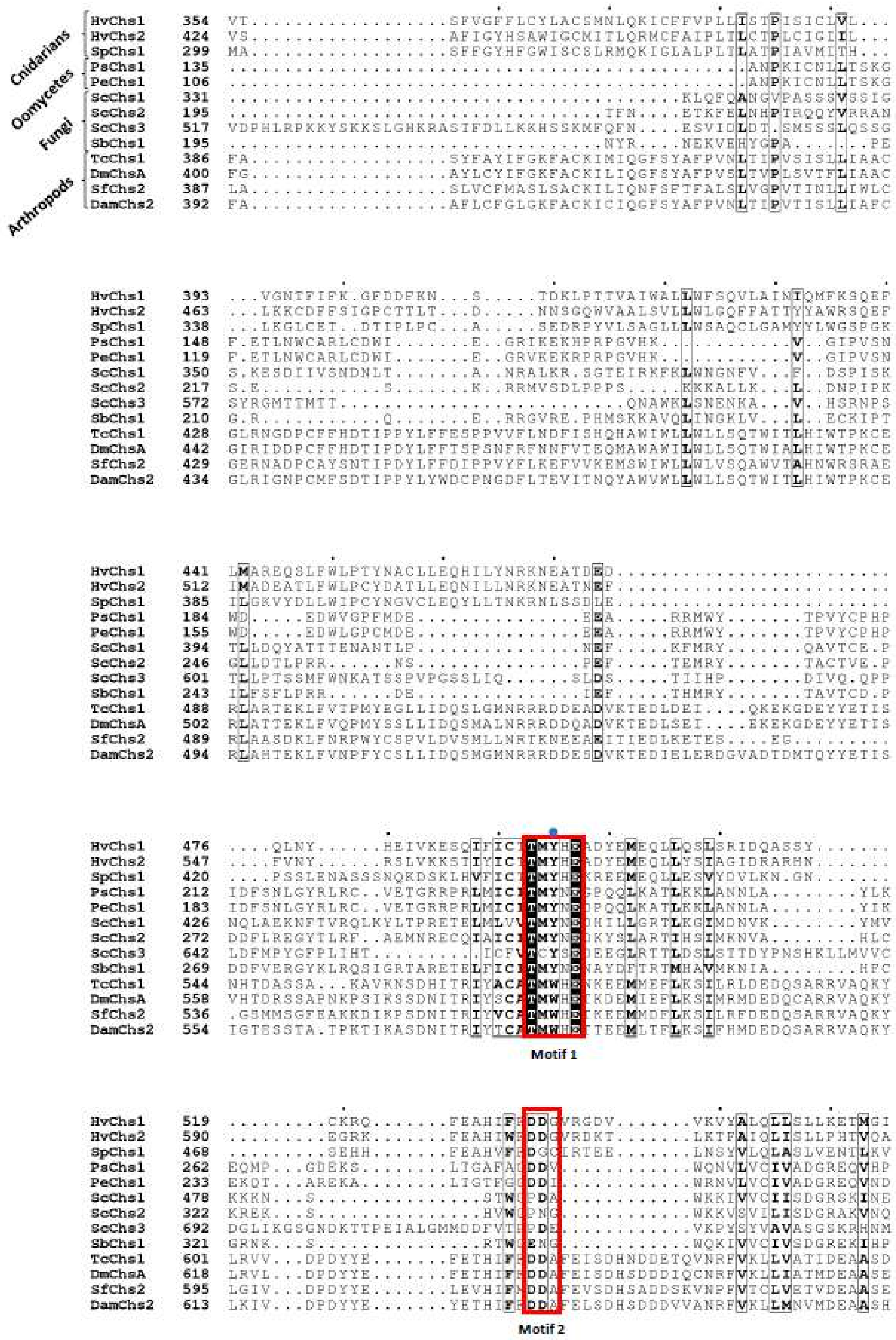
Cnidarians
Oomycetes
Fungi
Arthropods
Motif 1
Motif 2

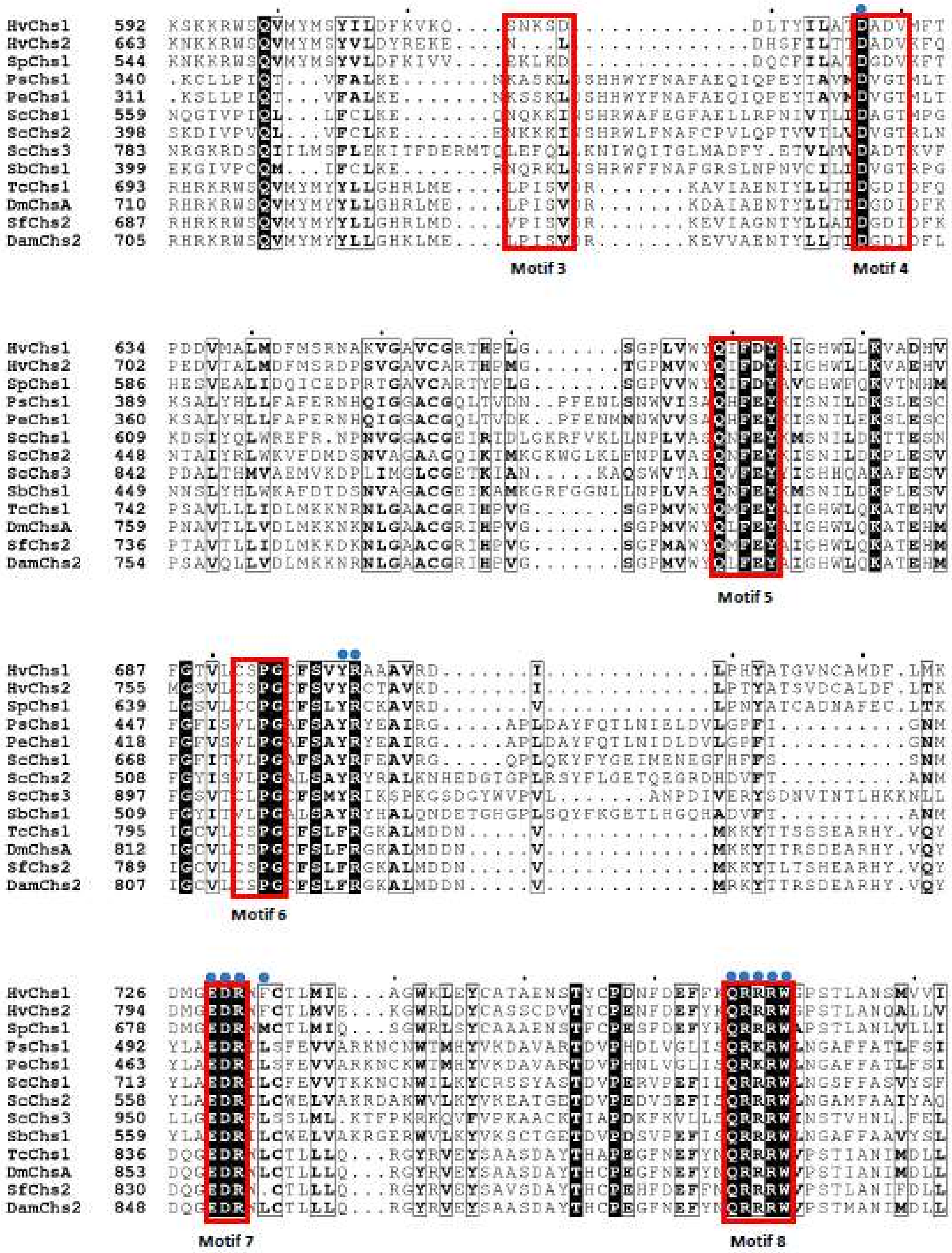

**Suppl. Fig. 2: Alignment of the two *Hydra vulgaris* chitin synthases to other well described chitin synthases.** Highlighted in red are conserved structural motifs and blue dots mark conserved catalytic active domains.

**Details of siRNA mediated knockdown**

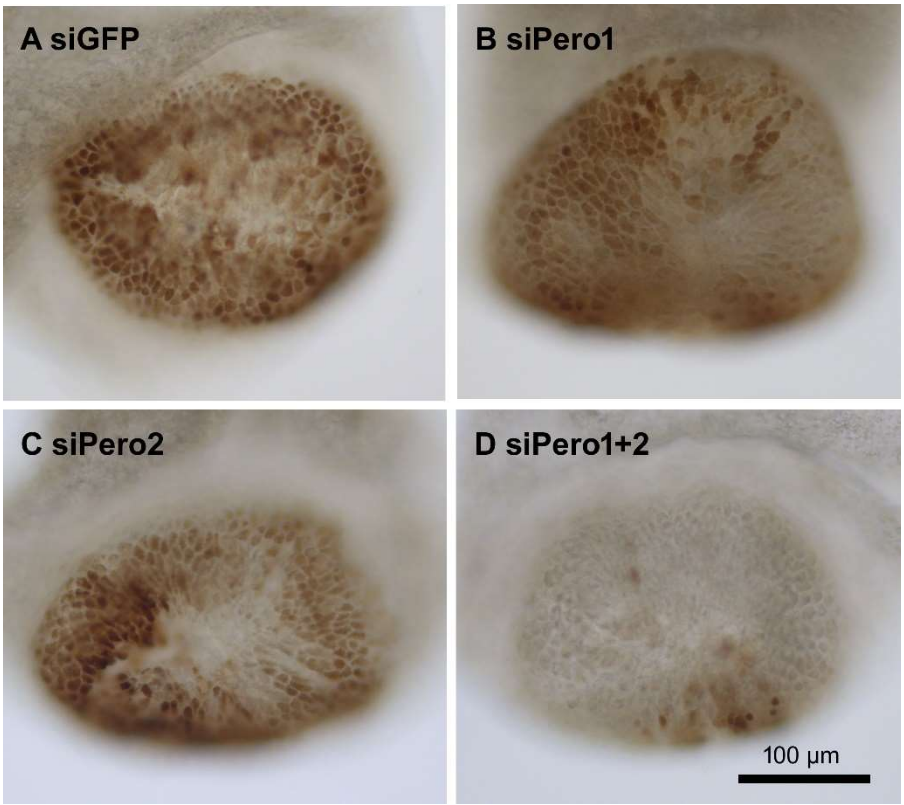


**Suppl. Fig. 3:** Peroxidase activity visualized with DAB staining in (A) siGFP (n=46), (B) Pero1 (n=21), (C) Pero2 (n=16), and (D) Pero1 + Pero2 double knockdown (n=26) basal discs. Note that single knockdown of Pero1 and Pero2 is not efficient to abolish Peroxidase staining.

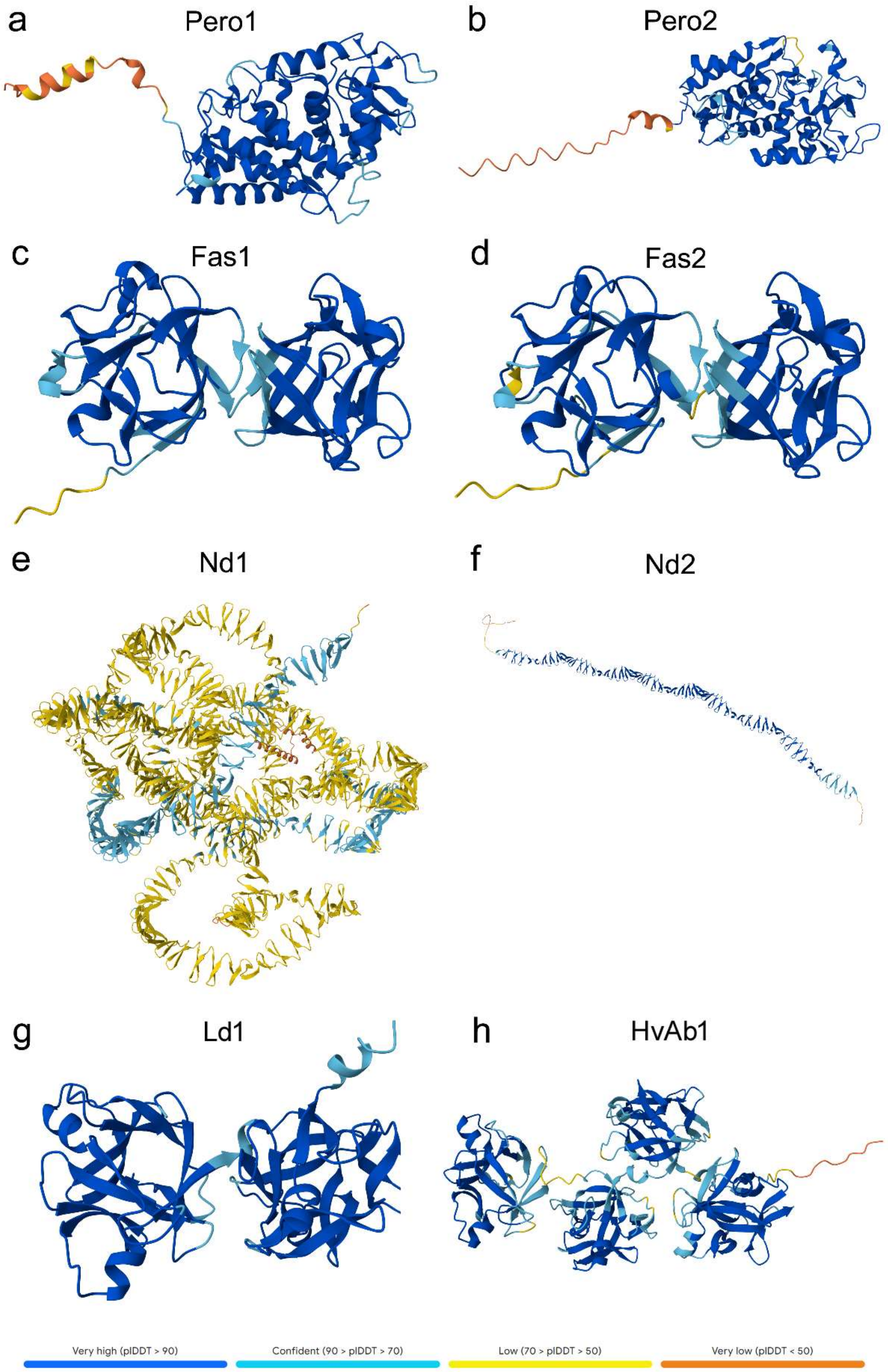


**Suppl. Fig.4: Predicted protein structures of siRNA target genes.** AlphaFold structure predictions of (a) Pero1, (b) Pero2, (c) Fas1, (d) Fas2, (e) Nd1, (f) Nd2, (g) Lb1, (h) HvAb1. Colours indicate confidence level of the predictions.

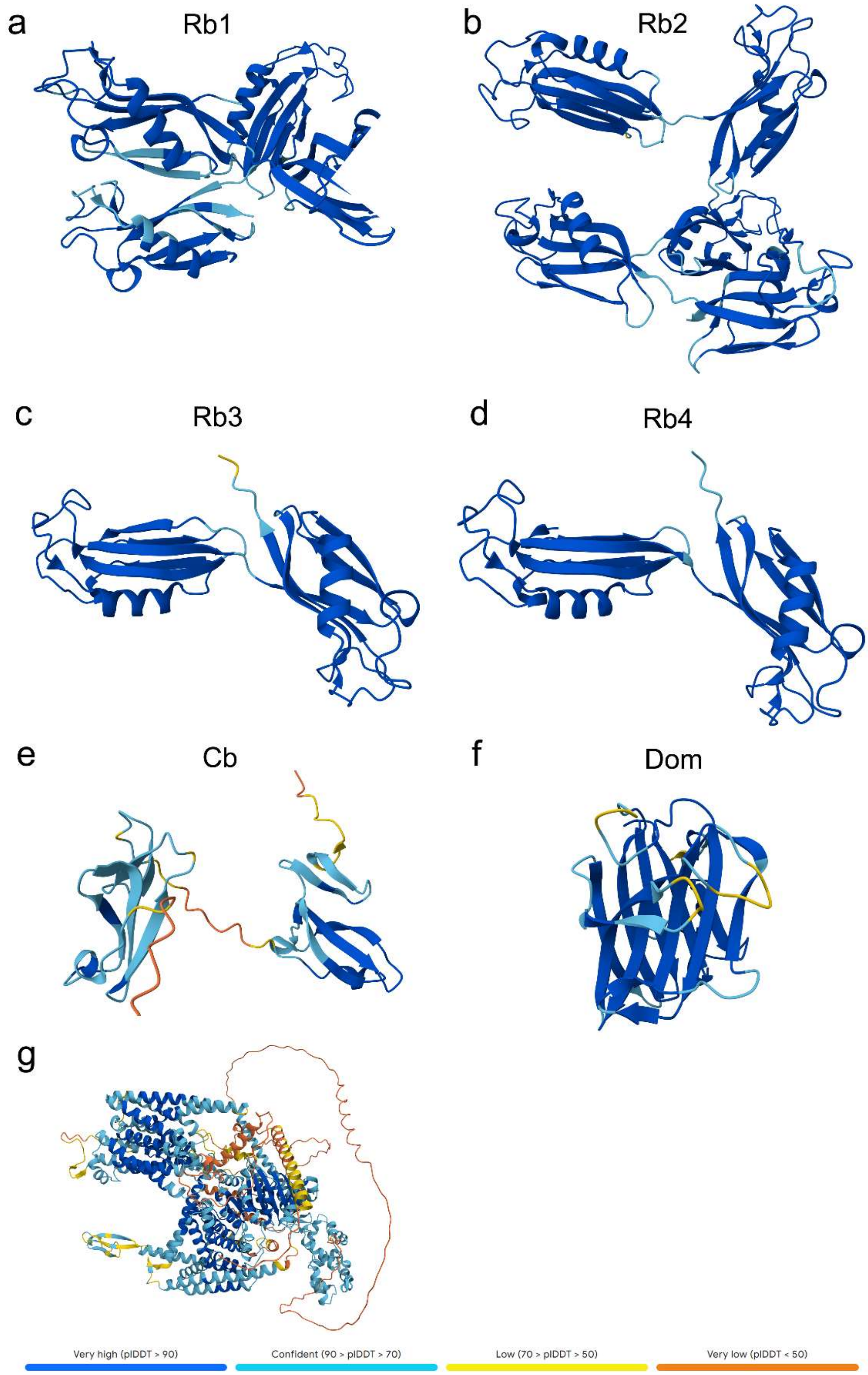


**Suppl. Fig.5: Predicted protein structures of siRNA target genes.** AlphaFold structure predictions of (a) Rb1, (b) Rb2, (c) Rb2, (d) Rb4, (e) Cb, (f) Dom, (g) Chs2. Colours indicate confidence level of the predictions.

**Details of siRNA mediated knockdown of multiple genes**

The siRNA sequences used for this publication are listed in Suppl. Table 1. The siRNAs were designed using the online tools siDirect v2.1 (Naito et al. 2009) and RNAxs (Tafer et al. 2008). Out of 20 predicted siRNA sequences, 14 resulted in a significantly reduced expression level of their target gene on mRNA level, indicating the high efficiency of these tools for siRNA prediction. For the double knockdown of Pero 1 and 2, and Lb1 and HvAb1, siRNAs targeting the single genes were used simultaneously at a concentration of 2 µM each. For double Fas1 and Fas2, Rb1 and Rb2, Rb3 and Rb4 knockdown, one siRNA targeting both sequences were used at an end concentration of 4 µM. The siRNAs designed for Nd1 and Nd2 both targeted both sequences and were used simultaneously at a concentration of 2 µM each. No separate knockdown of Fas1, Fas2, Nd1, Nd2, Rb1, Rb2, Rb3, and Rb4 was performed. For Dom knockdown, the combination of two siRNAs targeting the same gene showed to be more efficient than single siRNAs and two siRNAs were used simultaneously at a concentration of 2 µM each. For the knockdown of the five genes (Rb1, Rb2, Rb3, Rb4 and Lb1) and the six genes (Rb1, Rb2, Rb3, Rb4, Fas1 and Fas2) three siRNAs were used simultaneously at a concentration of 2 µM each.

Although survival rates did not differ significantly among treatments, they largely mirrored the experimenter’s level of experience. siPero1+2, siCb, and siDom treatments were conducted first, whereas siRb1+2, siRb3+4, siRb1+2+3+4+Lb1 and siRb1+2+3+4+Fas1+2 were conducted last. Over the time of the treatment, polyps started to bud and while freshly budded polyps shared the phenotype of the parental polys, only original polyps were further analysed.

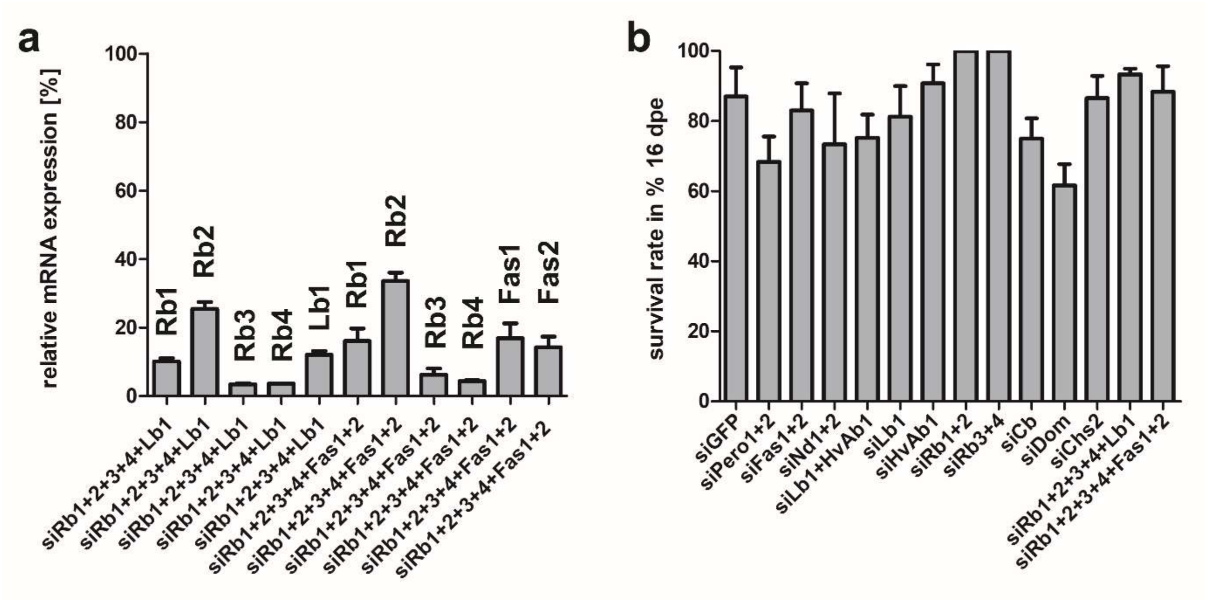


**Suppl. Fig. 6:** Knockdown efficiency after knockdown with three siRNAs and survival rate of knockdown treatment. (a) Knockdown efficiency in relative mRNA level after combining three siRNAs, qPCR target is indicated on top (means of relative mRNA expression level with SD, n=3). (b) Survival rate of knockdown procedure at 16 days past the first electroporation (16dpe), separated by siRNAs (means with SD, n≥3).

**Suppl. Table 1:** List of siRNA sequences.

| **Target** | **Si-Name** | **Sequence** | **notes** |
|---|---|---|---|
| Chs2 (HVAEP2.G004025) | siCHS2-3 | GAATGTACCGATGAAAATA | |
| Pero1 (HVAEP1-G001119) | siPero1-1 | GGATTTGACTTTAATTGCA | |
| Pero2 (HVAEP15-G028410 ) | siPero2-1 | GGCATTAGATGGATTTAAA | |
| Fas1 + Fas 2 (HVAEP1.G009194+ HVAEP1.G009216) | siFas-1 | GCGTGATTGCAAAAGAAAA | Double knockdown Fas 1 + Fas 2 |
| Nd1 (HVAEP3.G006596) | siND-A | GTTTTGTAACAACGATAAA | Double knockdown Nd1 and Nd2 |
| Nd2 (HVAEP3.G006609) | siND-B | GTATGTGGATCAAATGTTT | Double knockdown Nd1 and Nd2 |
| Lb1 (HVAEP1.G002182) | siAB1A | GTATTATCCTGGTTATGTT | |
| | siAB1B | GCATCGTTAGAGCATTAAA | |
| HvAb1 (HVAEP5.G010204) | siAB2B | GCTTGAATGGAATGATTAA | |
| Rb1 + Rb2 (HVAEP11.G020671+ HVAEP11.G020675 | siRb3-6 | GTAGAACATTGTCTAATGT | Double knockdown Rb1 + Rb2 |
| Rb3 + Rb4 (HVAEP11.G020720+ HVAEP11.G020722) | siRb68-6 | CATAAACTGTAATAACCAA | Double knockdown Rb3 + Rb4 |
| Cb (HVAEP8.G014104) | siCBP1-2 | CCTCAGAGTTTAATGTTTA | |

| | | |
|---|---|---|
| Dom (HVAEP3.G006789) | siDom-1 | CGAAATTTCAGAAATACAT |
| | siDom-2 | GTAATCGACTAGTTACAAA |

**Suppl. Table 2:** List of qPCR primers.

| **Target** | **Primer name** | **Sequence** |
|---|---|---|
| EF 1α | qPCR_EF1a_FW | TCAGGATGGCACGGAGATAATA |
| | qPCR_EF1a_RV | TGGTACAAAGGGTGGGAAGTAGA |
| RP 16 | RP16_qPCR_FW | TCTGTTGCTGAGGAGCGTGC |
| | RP16_qPCR_RV | AGTCCACGAAGTTCTCTGTGTTT |
| Chs2 (HVAEP2.G004025) | qPCR_CHS2_FW | TGCGAACGAGAAGCTGCTAT |
| | qPCR_CHS2_RV | TTCCAACGCCACCCCATAAT |
| | Pero-1_qPCR_FW | AGGCGAAGAACACTGTGATGG |
| Pero1 (HVAEP1-G001119) | Pero-1_qPCR_RV | TCGGCGGTGGATCTAGTTAATG |
| Pero2 (HVAEP15-G028410 ) | Pero-2_qPCR_FW | TCAGGCACTGTTCGTCTAGC |
| | Pero-2_qPCR_RV | AGTGAGAGCAACAACTGCGG |
| Fas1 (HVAEP1.G009194) | qPCR_Fasc1_FW | ACGAGATACACCTGTATGGACA |
| | qPCR_Fasc1_RV | TGTTTTTGCGTTAGTGAAACGA |
| Fas 2 (HVAEP1.G009216) | qPCR_Fasc2_FW | ACGAACTACACCTGTATGGACA |
| | qPCR_Fasc2_RV | TCCCATAAACCAATAGCATCTCT |
| Nd1 (HVAEP3.G006596) | qPCR_ND_A_FW | TATCAGTTGTGTGGAAGTGGC |
| | qPCR_ND_A_RV | AAGTTTGTACACTCCATCACC |
| Nd2 (HVAEP3.G006609) | qPCR_ND_B_FW | AACAAATGGAGCGATGAAAATGA |
| | qPCR_ND_B_RV | TACGTCGCTTCCACACATCT |
| Lb1 (HVAEP1.G002182) | qPCR_AB_1_FW | TGATGCTTCATTCATTATTTGCCCT |
| | qPCR_AB_1_RV | TCTTCTGAAAAGTTCTTCTTGAGGT |
| HvAb1 (HVAEP5.G010204) | qPCR_AB_2_FW | TGAAGCCGGCTGGAATAAGG |
| | qPCR_AB_2_RV | TAGAAGCGTCATCGGGGTTG |
| Rb1 (HVAEP11.G020671) | Rb3-qPCR-1_FW | TTTTGTCGGAGTTGCTCATGCC |
| | Rb3-qPCR-1_RV | GCAGTCGATTTTCAGTCCATACCC |
| Rb2 (HVAEP11.G020675) | Rb5-qPCR-1_FW | CGACGAACCAGATTCTCATGTTG |
| | Rb5-qPCR-1_RV | TTTGAATCGTAGGCACCAGGAC |
| Rb3 (HVAEP11.G020720) | Rb6-qPCR-1_FW | TGACGAACAAAAGGCAACCATAG |
| | Rb6-qPCR-1_RV | GTGATTGACCAGCAGCTCCAAA |
| Rb4 (HVAEP11.G020722) | Rb8-qPCR-2_FW | AGATGGCAAAAGGAGTAGTACCAC |
| | Rb8-qPCR-2_RV | CGAAGTTGTGATTGACCAGCAG |
| Cb (HVAEP8.G014104) | CBP_qPCR_FW | CAGTAGCACCTCCTTGCCC |
| | CBP_qPCR_RV | GTGTAACATGCAGTGTCTCCTTC |
| | qPCR_domon_FW | GTAAAACTAGCAGTTGTGATGTCTT |
| Dom (HVAEP3.G006789) | qPCR_domon_RV | CGCCCTTCATACCAGCCATT |

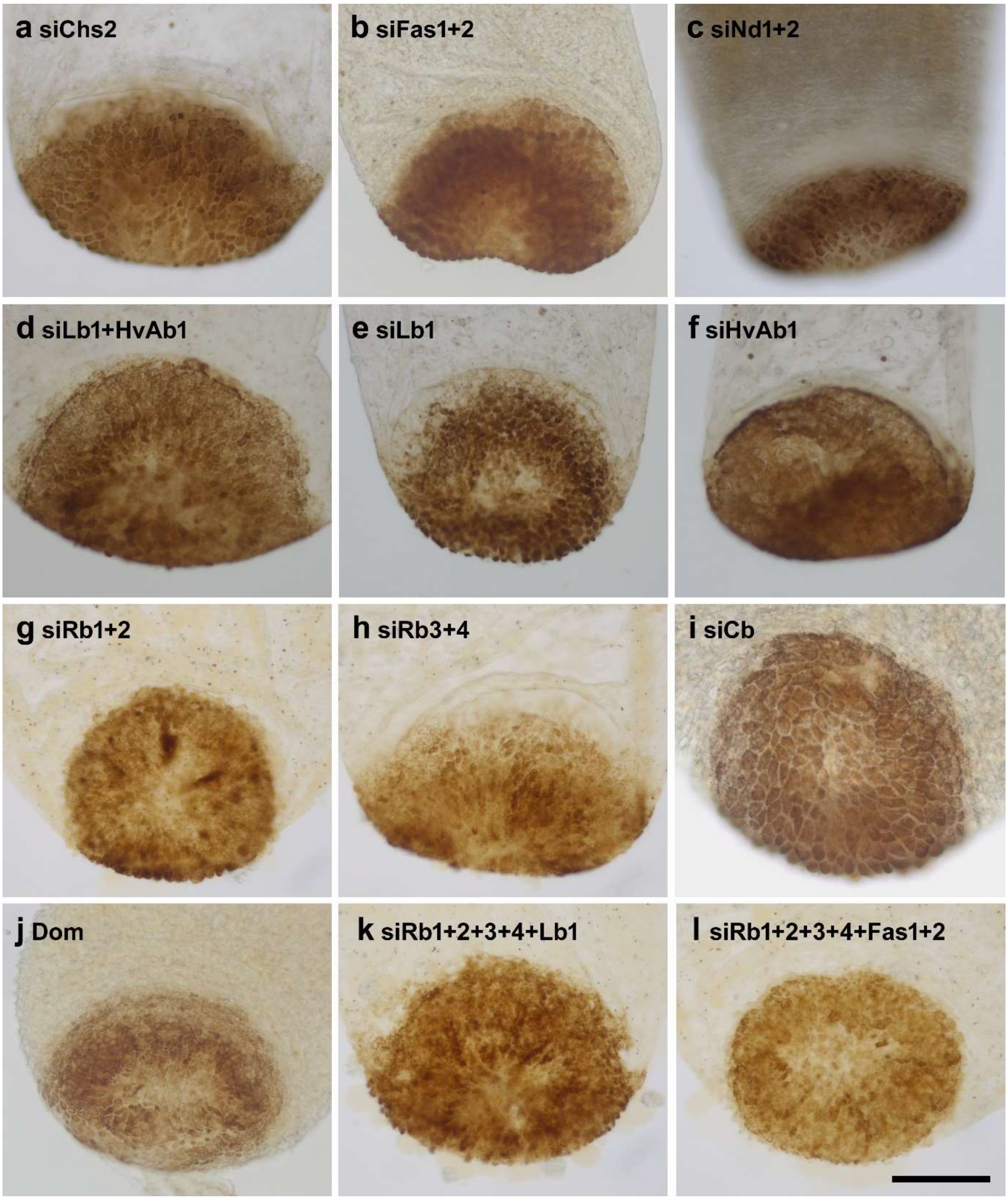


**Suppl. Fig. 7: Basal disc morphology and peroxidase activity in knockdown animals appeared normal.** Basal discs with peroxidase staining (DAB) after knockdown of (a) siChs2 (n = 17), (b) siFas1+2 (n=8), (c) siNd1+2 (n=9), (d) siLb1+2 (n = 20), (e) siLb1 (n = 15), (f) siHvAb1 (n = 16), (g) siRb1+2 (n=9), (h) Rb3+4 (n=10), (i) siCb (n=14), (j) siDom (n=10), (k) siRb1+2+3+4+Lb1 (n=10), (l) siRb1+2+3+4+Fas1+2 (n=10). Scale bar: 100 µm.

**Material and methods: TEM preparation**

For preparation, five Hydra polyps from each sample were relaxed in 2% urethane for 2 min to prevent stress-induced shrinkage of the peduncle. For siGFP and siHvAb1 samples chemical fixation was carried out using 1% glutaraldehyde and 0.2% $OsO_4$ in buffer, as described by Campbell et al. (1987). For siGFP and siChs samples chemical fixation was carried out with a 1:1 ratio of fixative A and fixative B (Shigenaka, Roth, and Pihlaja 1971). For all samples, dehydration was performed using a graded acetone series (2 × 50%, 2 × 70%, 90%, 3 × 100%). The polyps were then infiltrated and embedded in Embed-812 resin. Ultra-thin sections (80 nm) of Hydra were cut longitudinally using an Ultracut UTC (Leica, Austria) and stained for two minutes with Reynolds stain (Reynolds, 1963). Ultra-thin section images of Hydra vulgaris longitudinal sections were visualized with a Libra 120 energy filter transmission electron microscope (Zeiss, Germany). They were obtained via the TRS 2x2k high-speed camera (Tröndle, Germany) combined with the ImageSP software (SYSPROG, Belarus). The images were edited using Photoshop (Adobe, Ireland).

**Morphology of basal discs remained unaltered by HvAb1 knockdown**

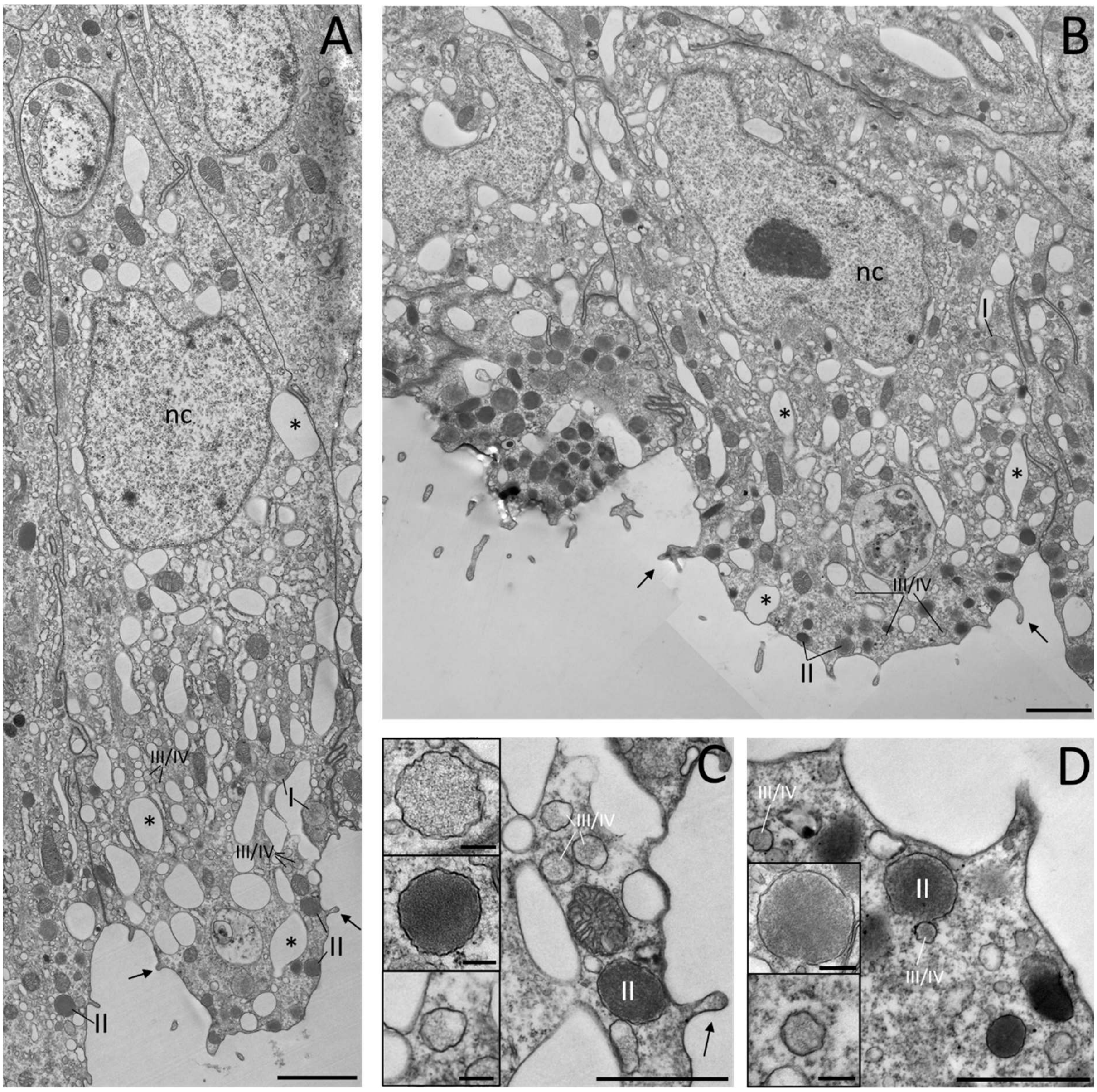


**Suppl. Fig. 8: Ultrastructure of basal disc cells of siGFP (control) and siHvAb1 knockdown animals.** (A-B) Longitudinal sections through basal disc showing individual basal disc cells from (a) siGFP, (b) siHvAb1 knock-down polyps. (c-d) Detail images of basal disc cells highlighting the different types of secretory granules (c) GFP knockdown, (d) HvAb1 knockdown. Insets show representative adhesive granules. Scale bars: (a-b) 2 µm, (c-d) 1 µm, (Insets) 150 nm. Abbreviations: nc=nucleus, I=Hydra secretory granule I, II=Hydra secretory granule II, III/IV=Hydra secretory granule III or IV, arrows indicate cytoplasmic extensions, asterisks indicate vacuoles of water.

a

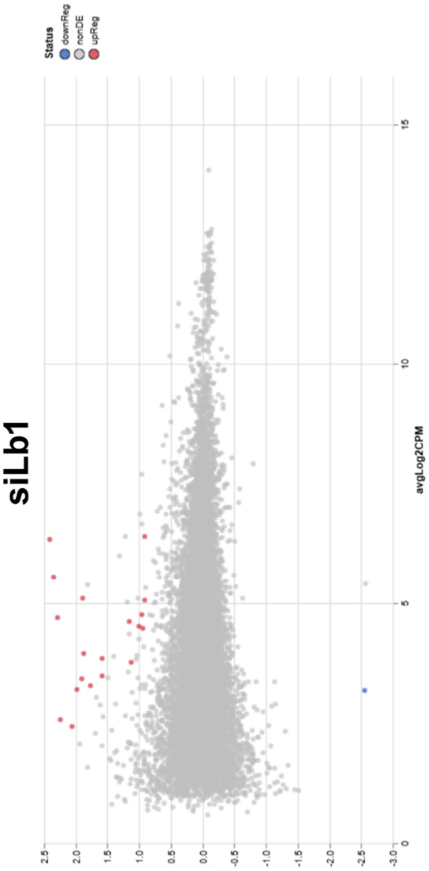


| full_gene_id | logFC | logCPM | F | PValue | FDR |
|---|---|---|---|---|---|
| HVAEP12.G023165 | 2,345984 | 5,567412 | 90,04027 | 6,54E-08 | 0,000762 |
| HVAEP13.G024463 | 2,060005 | 2,441342 | 58,65788 | 3,79E-07 | 0,00148 |
| HVAEP12.G023132 | 2,414335 | 6,356542 | 68,92757 | 3,81E-07 | 0,00148 |
| HVAEP11.G020918 | 1,910454 | 3,433744 | 55,96739 | 1,46E-06 | 0,003407 |
| HVAEP13.G025703 | 1,982354 | 3,220801 | 48,91006 | 1,83E-06 | 0,003407 |
| HVAEP12.G021832 | 0,918468 | 6,422945 | 53,45838 | 1,91E-06 | 0,003407 |
| HVAEP2.G004844 | 1,764103 | 3,306886 | 51,84766 | 2,31E-06 | 0,003407 |
| HVAEP11.G020700 | 1,152123 | 4,62968 | 51,28136 | 2,47E-06 | 0,003407 |
| HVAEP9.G015809 | 1,580016 | 3,505994 | 50,72354 | 2,63E-06 | 0,003407 |
| HVAEP2.G004757 | 1,581762 | 3,873995 | 47,57869 | 3,9E-06 | 0,004543 |
| HVAEP3.G006016 | 0,961701 | 4,782125 | 36,92516 | 1,45E-05 | 0,014664 |
| HVAEP10.G018266 | 2,245122 | 2,59478 | 37,86528 | 1,51E-05 | 0,014664 |
| HVAEP6.G010754 | 0,905551 | 5,075674 | 35,83588 | 1,99E-05 | 0,016828 |
| HVAEP2.G004733 | 1,880134 | 3,960941 | 35,3333 | 2,19E-05 | 0,016828 |
| HVAEP6.G011105 | 2,278618 | 4,720476 | 34,69908 | 2,42E-05 | 0,016828 |
| HVAEP6.G011135 | 1,891845 | 5,126429 | 34,64685 | 2,44E-05 | 0,016828 |
| **HVAEP1.G002182 (=Lb1)** | **-2,55038** | **3,189771** | **34,67102** | **2,46E-05** | **0,016828** |
| HVAEP3.G006245 | 0,941054 | 4,496756 | 32,34518 | 3,57E-05 | 0,023068 |
| HVAEP9.G017177 | 0,995456 | 4,526916 | 30,78876 | 4,65E-05 | 0,028521 |
| HVAEP3.G005995 | 1,127755 | 3,777326 | 28,67012 | 6,84E-05 | 0,039807 |

b

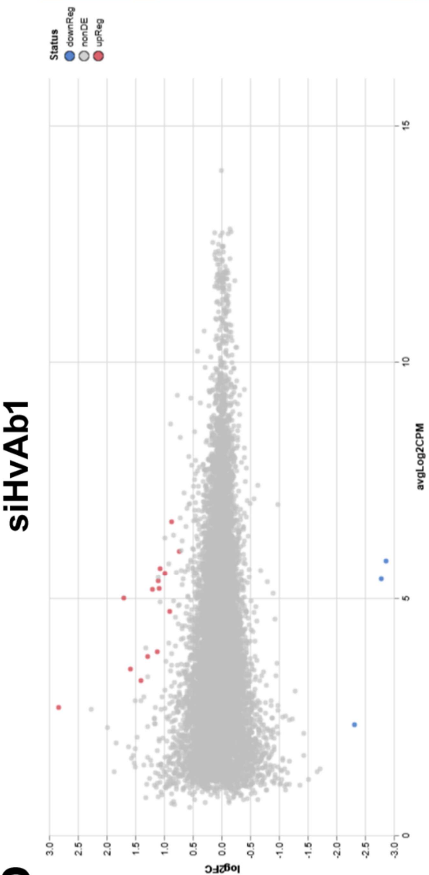


| full_gene_id | logFC | logCPM | F | PValue | FDR |
|---|---|---|---|---|---|
| **HVAEP5.G010204 (=HvAb1)** | **-2,86144** | **5,816902** | **138,926** | **3,17E-09** | **3,69E-05** |
| HVAEP14.G026829 | 2,83201 | 2,714239 | 50,27115 | 2,04E-06 | 0,011849 |
| HVAEP8.G014559 | 0,861312 | 6,647389 | 44,08321 | 6,14E-06 | 0,018066 |
| HVAEP7.G013564 | 1,061499 | 5,647625 | 43,61658 | 6,54E-06 | 0,018066 |
| HVAEP8.G015302 | -2,31913 | 2,34318 | 48,01055 | 7,76E-06 | 0,018066 |
| HVAEP1.G000761 | 1,094182 | 5,388137 | 40,98324 | 9,57E-06 | 0,018545 |
| HVAEP12.G024012 | 1,114541 | 3,885059 | 39,80769 | 1,11E-05 | 0,018545 |
| HVAEP3.G005995 | 1,276667 | 3,777326 | 37,27349 | 1,64E-05 | 0,023837 |
| HVAEP1.G001434 | 1,082653 | 5,224461 | 34,998 | 2,31E-05 | 0,027477 |
| HVAEP15.G027635 | 1,702473 | 5,023394 | 34,65801 | 2,44E-05 | 0,027477 |
| HVAEP6.G011284 | 0,72663 | 6,002841 | 34,27252 | 2,60E-05 | 0,027477 |
| HVAEP12.G021405 | 0,986118 | 5,540059 | 33,47774 | 2,96E-05 | 0,028692 |
| HVAEP4.G008584 | 1,396037 | 3,285418 | 32,35357 | 3,57E-05 | 0,031987 |
| HVAEP2.G005042 | 1,575878 | 3,526234 | 30,32423 | 5,07E-05 | 0,040252 |
| HVAEP7.G012476 | 1,197689 | 5,192908 | 30,17051 | 5,19E-05 | 0,040252 |
| HVAEP10.G018629 | -2,77718 | 5,419949 | 29,72695 | 5,61E-05 | 0,04083 |
| HVAEP12.G023182 | 0,896046 | 4,73132 | 28,57265 | 6,91E-05 | 0,047329 |

**Suppl. Fig. 9: Volcano blot indicating differential expressed genes between siGFP, siLb1, and siHvAb1 knockdown samples with a false discovery rate (FDR) <0.05.** (a) Differentially expressed genes between siLb1 and siGFP, volcano blot and list of genes. (b) Differentially expressed genes between siHvAb1 and siGFP, volcano blot and list of genes. Red dots indicate upregulated and blue dots downregulated genes (n=3).

**RNA-seq gene expression analysis**

Total RNA of a cohort of 10 siRNA knockdown animals (siGFP, siLb1, and siHvAb1) at 17 dpe were isolated using the Monarch® Total Miniprep Kit (New England BioLabs) according to the manufacturers protocol. Total RNA of three biological samples was send to sequencing. Library preparation, sequencing and gene expression analysis were performed at the company XPseq Analytics GmbH (Austria) using their ExpressoSeq rapid 3' RNA-Seq service. Sequencing was performed on a PromethION 2 Solo sequencer on a R10 flow cell and the Ligation Sequencing Kit V14 (Oxford Nanopore Technologies). Sequencing reads were aligned to the *Hydra vulgaris* AEP (PMID: 36639202) reference genome using the Minimap2 aligner (version 2.28). Read counting and differential gene expression analysis was performed with the Rsubread (version 2.22.1) and edgeR (version 4.6.3) packages in R (version 4.5.0).

Restricting read counting and differential gene expression analysis to basal-disc-specific genes yielded four differently expressed genes with a false discovery rate (FDR) of <0.05 in HvAb1 (aside from the siRNA target HvAb1) (see list below). No significantly different expressed basal-disc-specific gene (aside from the downregulated Lb1) was identified after knockdown of Lb1.

| full_gene_id | logFC_siLb1 | logFC_siHvAb1 | uncorrected_ PValue_siLb1 | uncorrected_ PValue_siHvAb1 | FDR_siLb1 | FDR_siHvAb1 |
|---|---|---|---|---|---|---|
| **HVAEP5.G010204 = HvAb1** | **0,330453331** | **-2,861435411** | **0,106864316** | **3,16763E-09** | **0,5754232** | **6,65202E-07** |
| HVAEP11.G020722 = Rb4 | 0,554679161 | 1,105518037 | 0,054146682 | 0,000655746 | 0,499237 | 0,037426738 |
| HVAEP2.G004393 | -0,368673996 | -0,793899155 | 0,059340737 | 0,000745364 | 0,499237 | 0,037426738 |
| HVAEP9.G016823 | 0,331362332 | 0,693521519 | 0,064952134 | 0,000681253 | 0,5246134 | 0,037426738 |
| HVAEP7.G012888 | 0,040658019 | 0,532382832 | 0,762677768 | 0,000891113 | 0,9590559 | 0,037426738 |

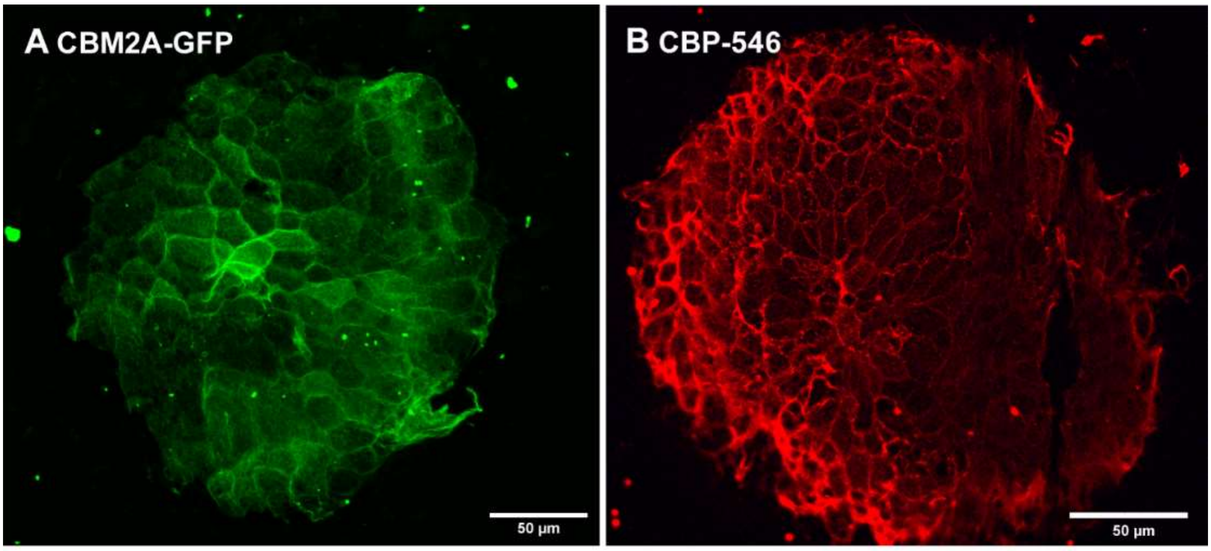


**Suppl. Fig. 10:** Chitin labelling of Hydra footprints with (A) CBM2A-GFP and (B) CBP-546, indicating the presence of chitin.

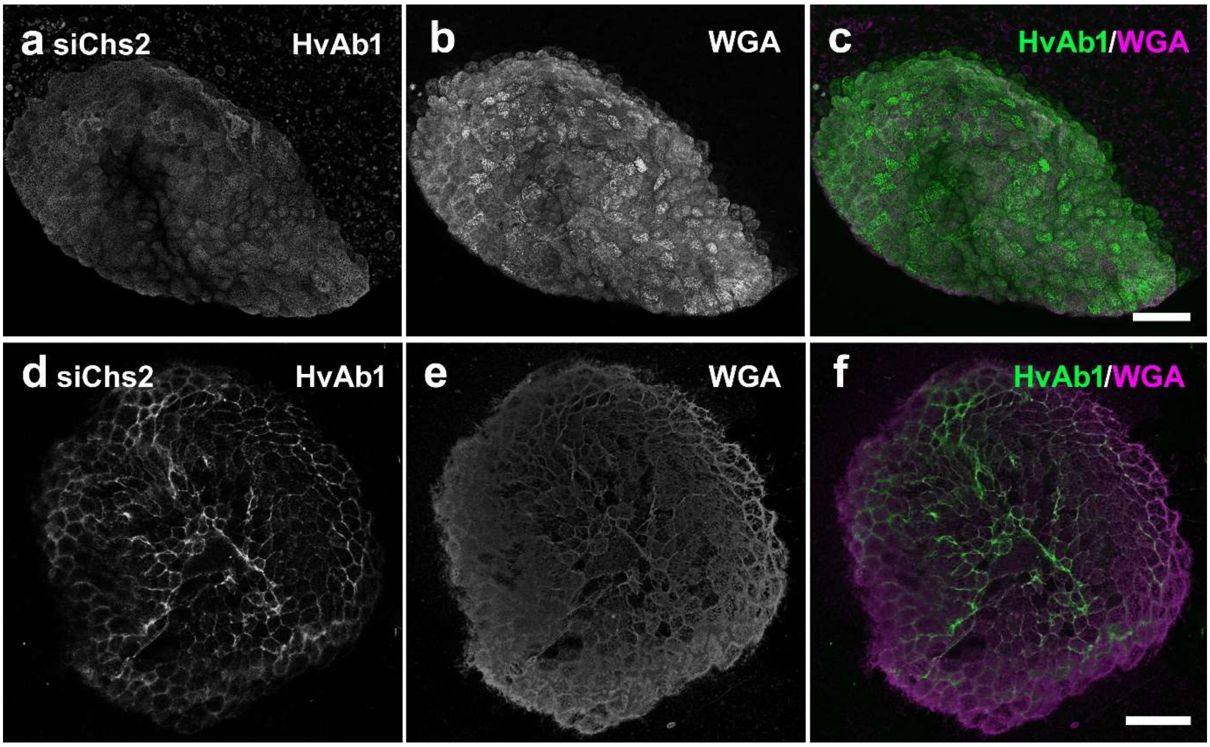


**Suppl. Fig. 11: HvAb1 and WGA distribution in Hydra basal disc and footprints after siChs2 treatment**. (a-c) Co-labelling of a siChs2 treated basal disc with (a) HvAb1, (b) WGA, and (c) the merged image. (d-f) Secreted footprint of a siChs2 treated animal showing (d) HvAb1, (e) WGA, and (f) the merged image (n=16). Scale bar: 50 µm.

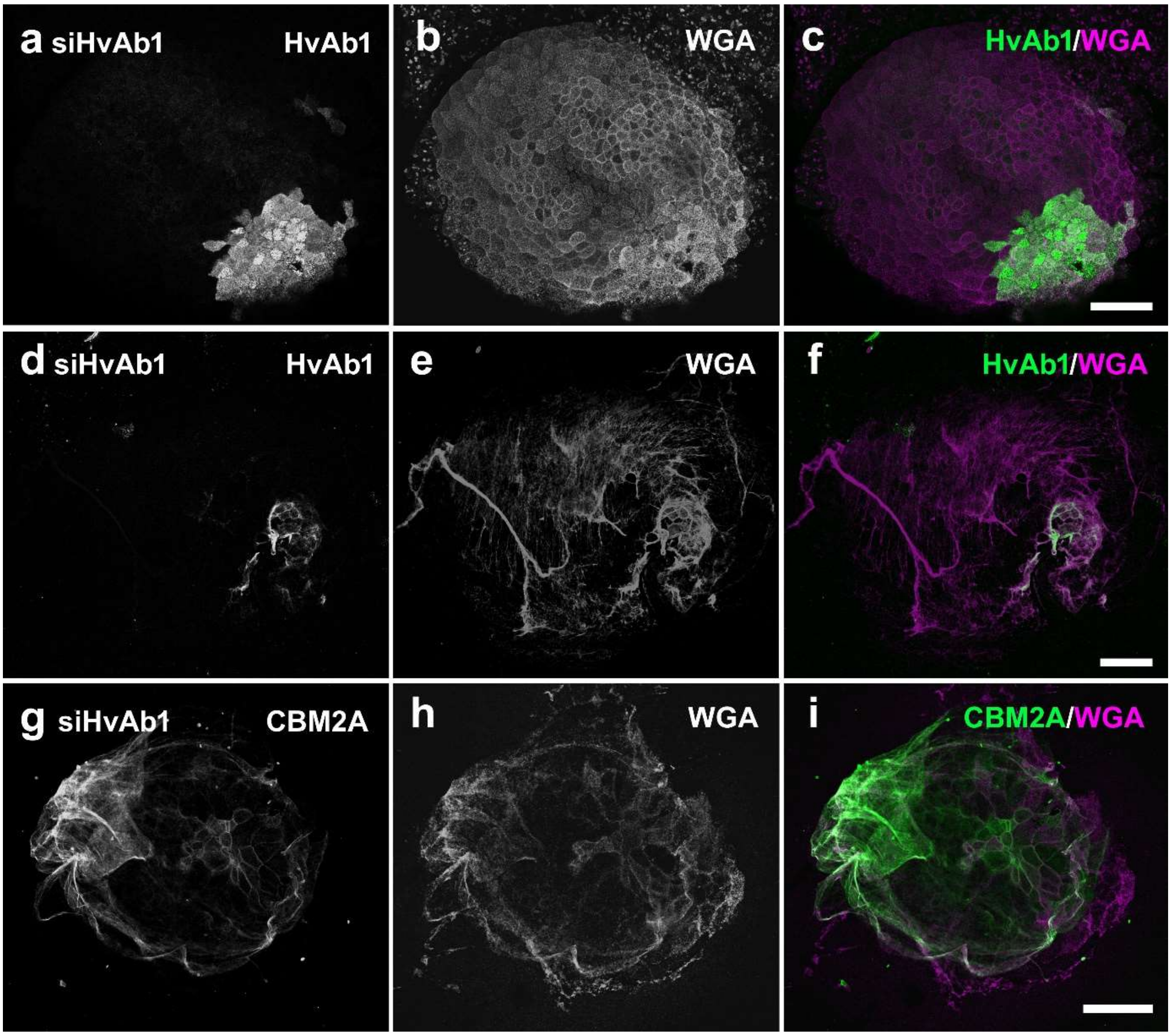


**Suppl. Fig. 12: Hydra basal disc and HvAb1, WGA, and CBM2A signals of corresponding footprints after siHvAb1 treatment.** (a-c) Co-labelling of a siHvAb1 treated basal disc with (a) HvAb1, (b) WGA, and (c) the merged image. (d-f) overview of the corresponding secreted footprint showing (d) HvAb1, (e) WGA, and (f) the merged image. (g-i) overview of the corresponding secreted footprint showing (d) CBM2A-GFP, (e) WGA, and (f) the merged image (n=21). Scale bars: 50 µm.

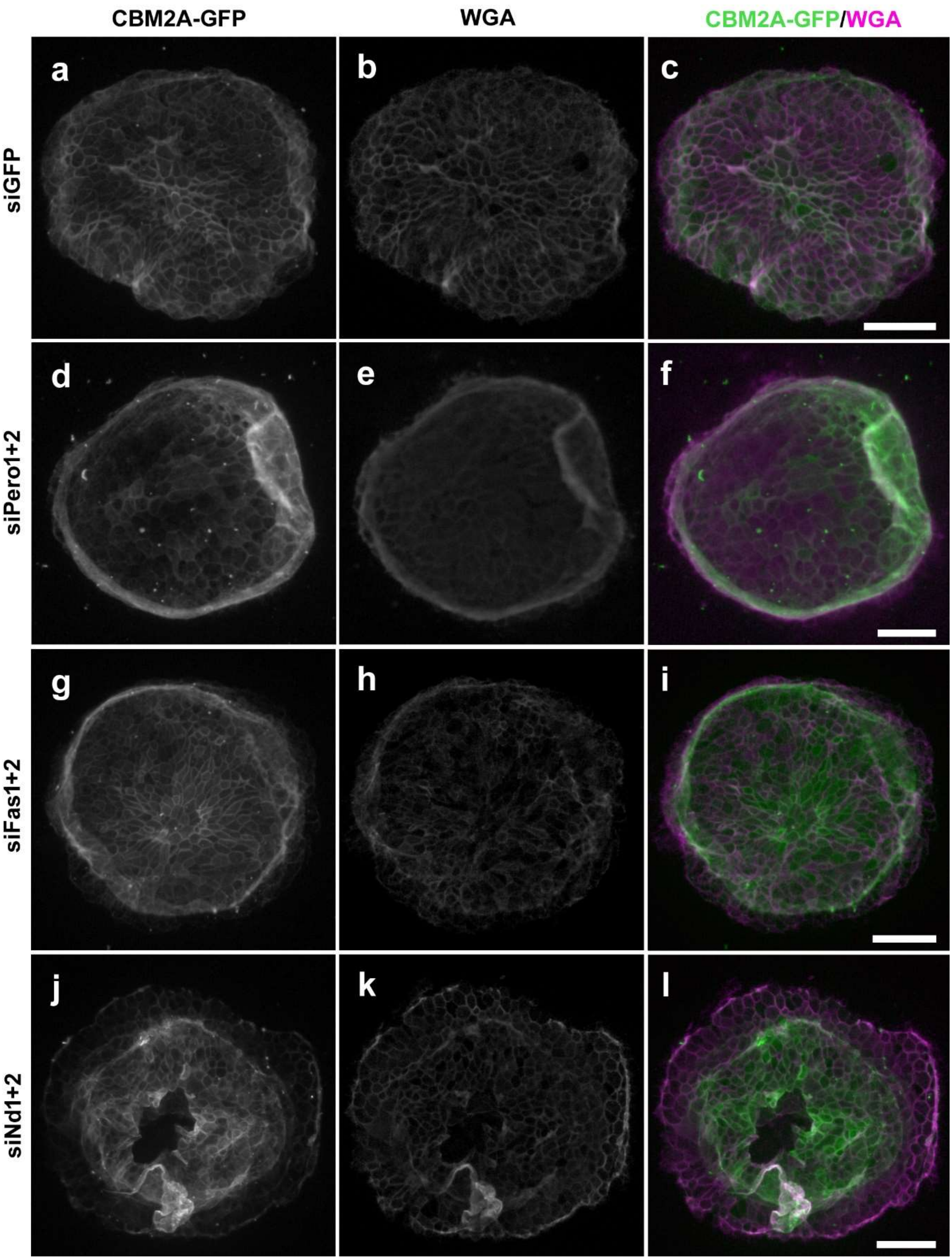


**Suppl. Fig. 13: Footprint structure visualized with CBM2A-GFP (chitin) and WGA staining of knockdowns that did not show any alterations.** CBM2A-GFP and WGA staining after knockdown of (a-c) siGFP, (d-f) siPero1+2, (g-i) siFas1+2, (j-l) siDom. Scale bars 50 µm.

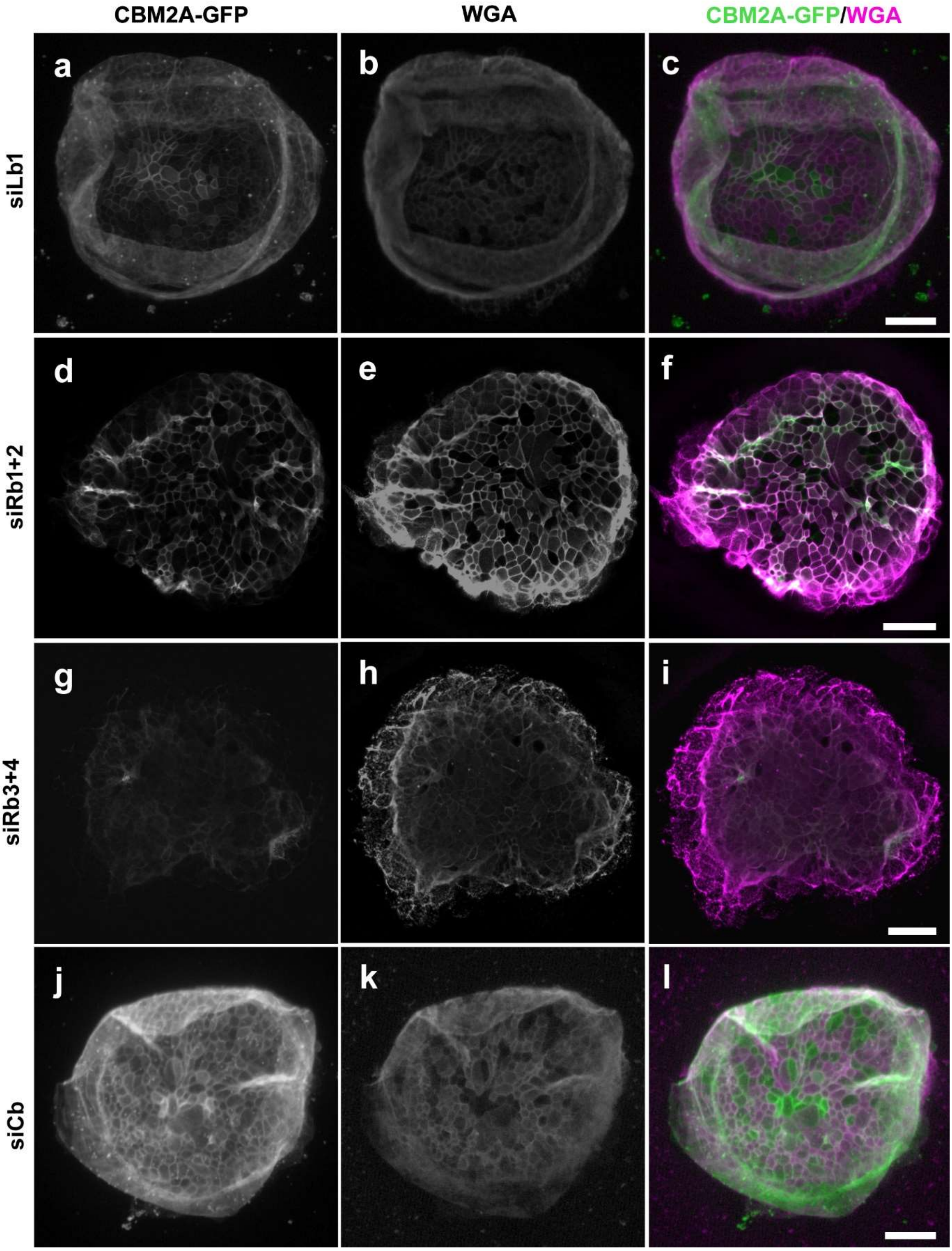


**Suppl. Fig. 14: Footprint structure visualized with CBM2A-GFP (chitin) and WGA staining of knockdowns that did not show any alterations.** CBM2A-GFP and WGA staining after knockdown of (a-c) siGFP, (d-f) siPero1+2, (g-i) siFas1+2, (j-l) siDom. Scale bars 50 µm.

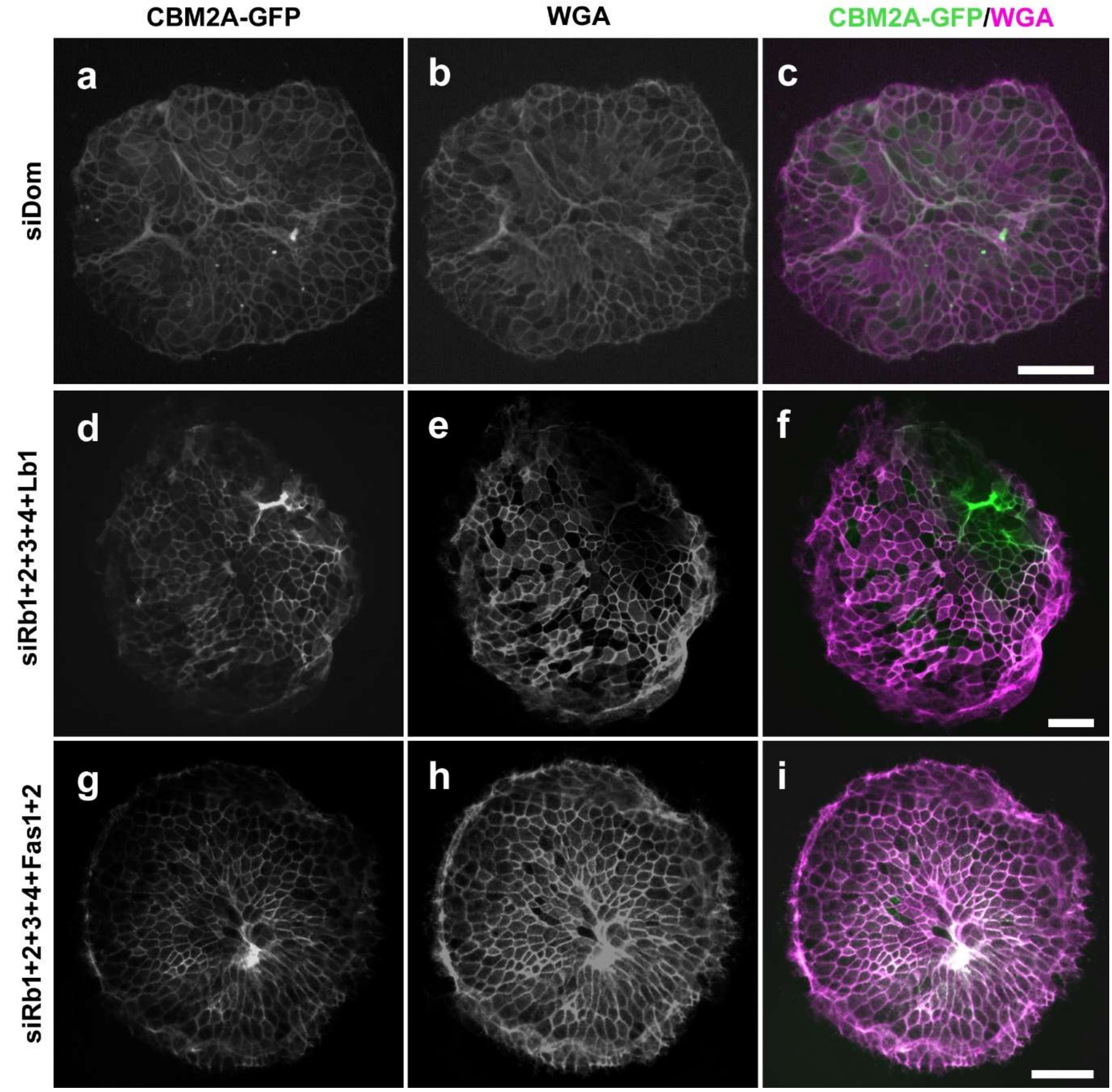


**Suppl. Fig. 15: Footprint structure visualized with CBM2A-GFP (chitin) and WGA staining of knockdowns that did not show any alterations.** CBM2A-GFP and WGA staining after knockdown of (a-c) siDom, (d-f) siRb1+2+3+4+Fas1+2, (g-i) siRb1+2+3+4+Lb1, (j-l). Scale bars 50 µm.

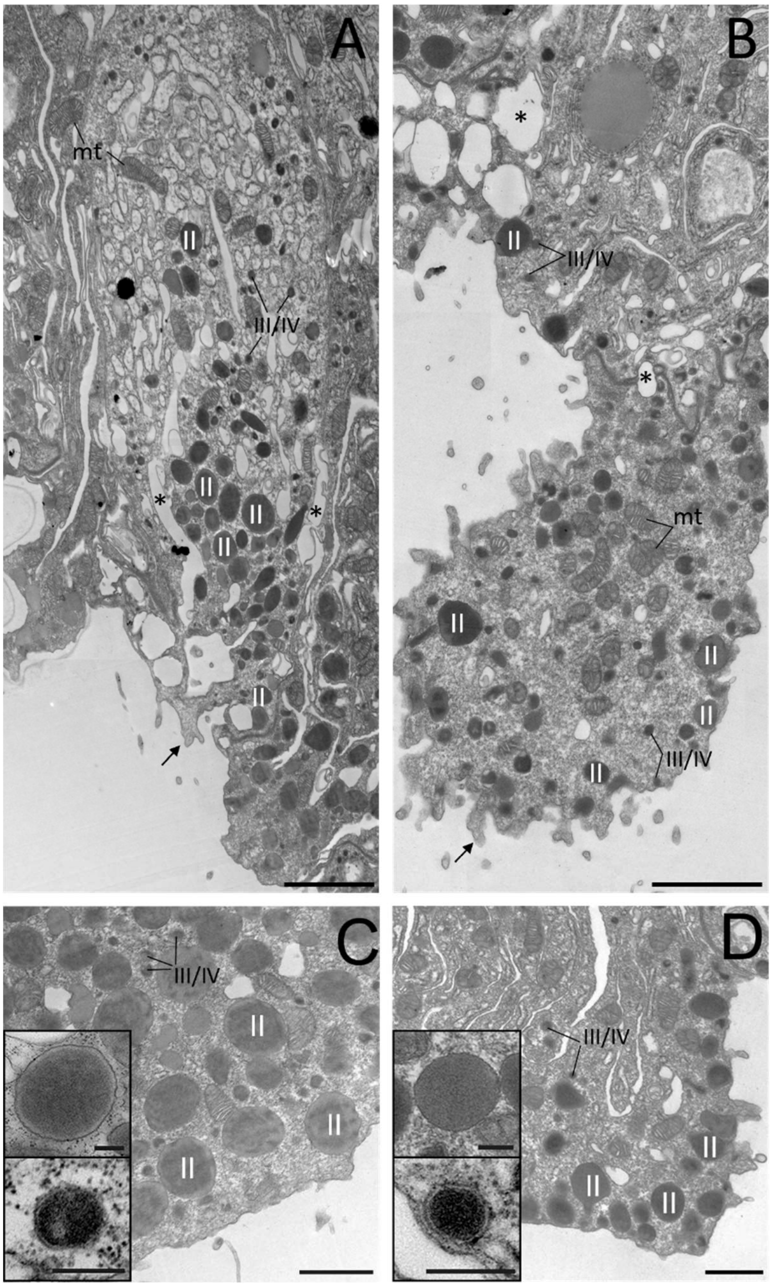


**Suppl. Fig. 16: Ultrastructure of basal disc cells after of control (siGFP) and siChs2 knockdown animals.** (a-b) Longitudinal sections through basal disc of (a) siGFP and (b) siChs2 knockdown animals. (c,d) Detail images of basal disc cells highlighting the different types of adhesive granules (c) siGFP, (d) siChs2 Insets show representative adhesive granules. Scale bars: (a,b) 2 µm, (c,d) 1 µm, (Insets) 250 nm. Abbreviations: mt=mitochondria, I=Hydra secretory granule I, II=Hydra secretory granule II, III/IV=Hydra secretory granule III or IV, arrows indicate cytoplasmic extensions, asterisks indicate vacuoles of water.

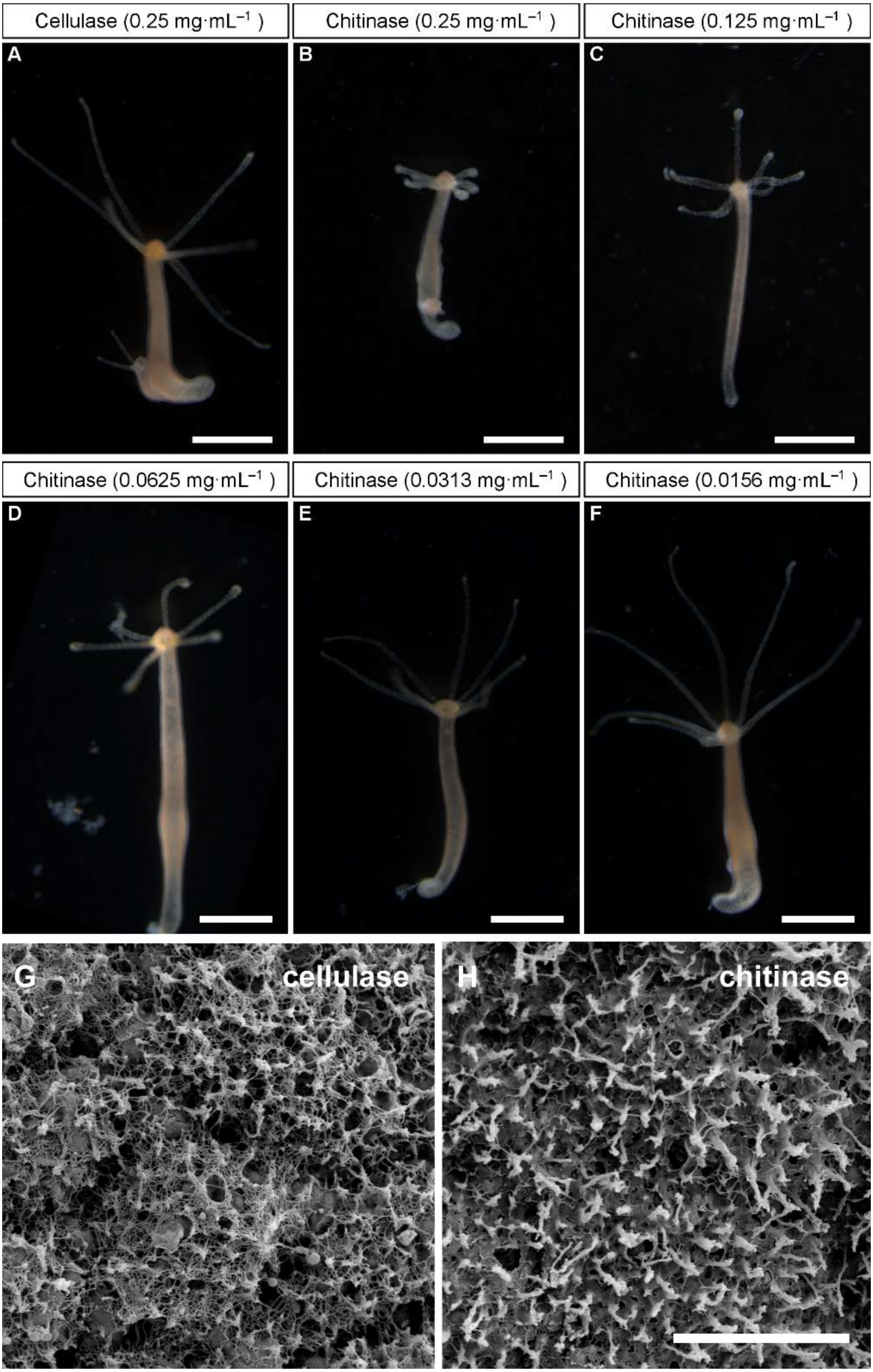
Cellulase (0.25 mg·mL⁻¹ )
Chitinase (0.25 mg·mL⁻¹ )
Chitinase (0.125 mg·mL⁻¹ )
A
B
C
Chitinase (0.0625 mg·mL⁻¹ )
Chitinase (0.0313 mg·mL⁻¹ )
Chitinase (0.0156 mg·mL⁻¹ )
D
E
F
G
cellulase
H
chitinase

**Suppl. Fig. 17: Whole mount polyps after the treatment with cellulase (control) and chitinase.** (A-F) Whole mount animals after treatment with decreasing concentrations of cellulase and chitinase (A) Cellulase treatment animals looked like untreated animals at all concentrations. (B) Highest chitinase treatment caused some shortening of tentacles, indicating some toxicity of the treatment, but animals remained viable and moved normally. (G-H) Scanning electron microscopy of the basal disc shows the (G) fibrous meshwork after cellulase (n=9), (H) but not after chitinase treatment (n=10). Scale bars (A-F) 100 µm, (G,H) 5 µm.

**SEM preparation of polyps**

Polyps were relaxed in 2 % urethane in *Hydra* medium and fixed in 4 % PFA in PBS overnight. Then the samples were washed in an ethanol series using 70%, 90% and 100% ethanol for at least 30 min each. Samples were critical point dried (BAL-TEC CPD 030) and polyps placed upside down on sample holders. The samples were then coated with 20 nm of a mixture of 80 % gold and 20 % palladium. A Zeiss DSM 982 GEMINI scanning electron microscope was used for the observation of the probes.

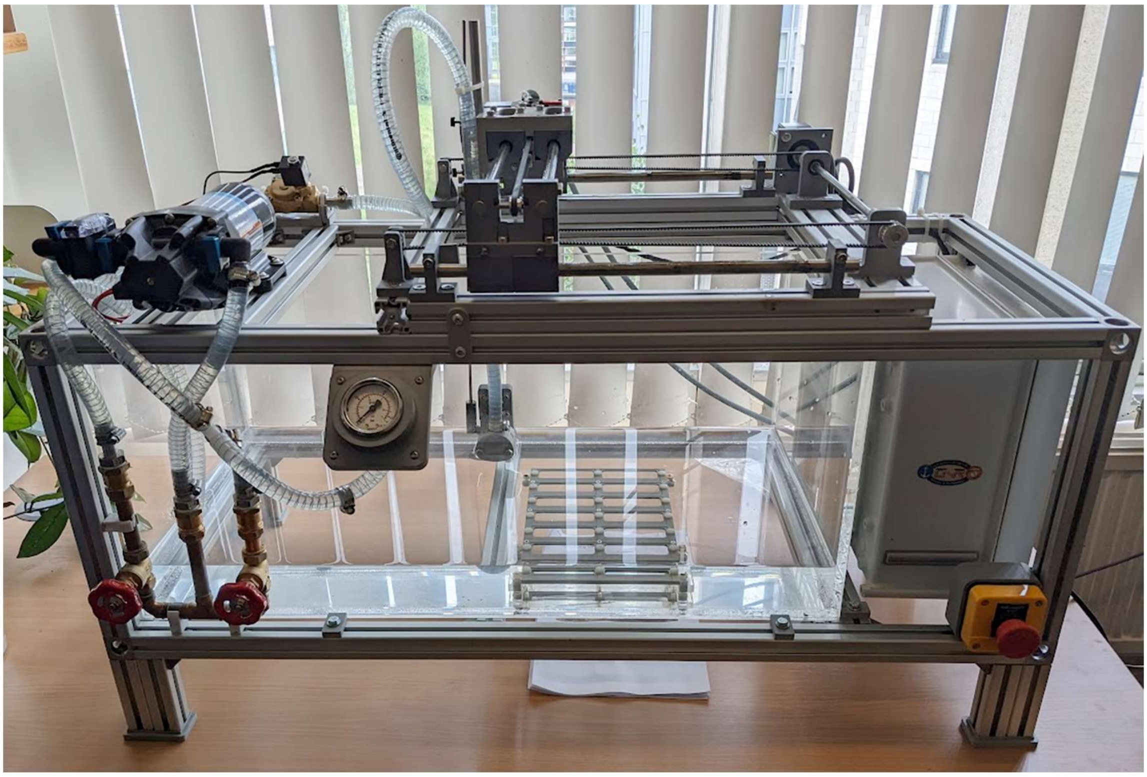

**Suppl. Fig. 18: Image of the water jet system.**

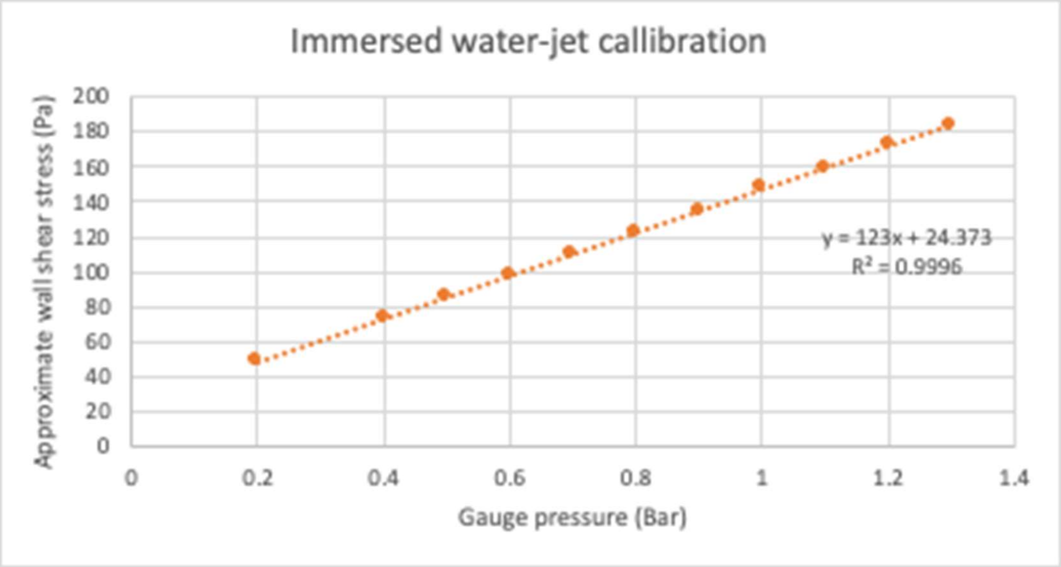


**Suppl. Fig. 19: The relationship between measured gauge pressure and approximate wall shear stress, calculated using equation described in material and methods.**

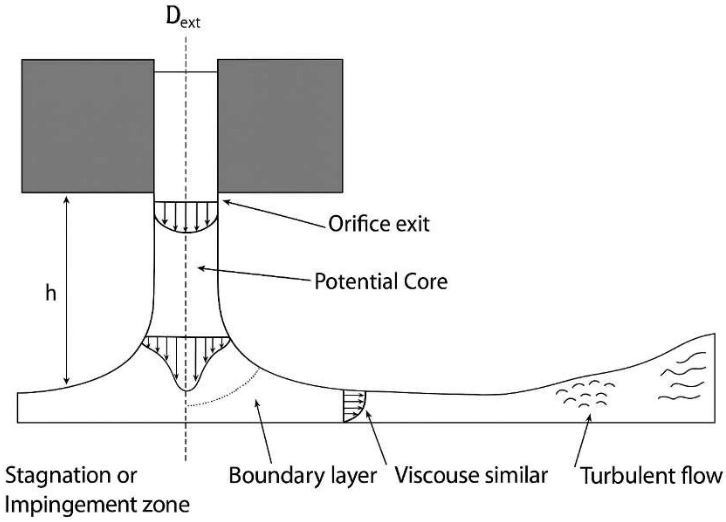


**Suppl. Fig. 20: : Schematic of the hydrodynamic regime discussed here.**
